# Efficient Identification of Short Tandem Repeats via Context-Aware Motif Discovery and Ultra-Fast Sequence Alignment

**DOI:** 10.1101/2025.11.25.690584

**Authors:** Xingyu Liao, Long Wen, Minghui Jing, Xingyi Li, Bolin Chen, Bin Zhang, Xin Gao, Xuequn Shang

**Affiliations:** School of Computer Science, Northwestern Polytechnical University (NPU), Xi’an, Shaanxi 710072, P.R. China.; Mohamed bin Zayed University of Artificial Intelligence, Masdar City, Abu Dhabi, United Arab Emirates; Computer Science Program, Computer, Electrical and Mathematical Sciences and Engineering Division, King Abdullah University of Science and Technology (KAUST), Thuwal 23955-6900, Kingdom of Saudi Arabia; Center of Excellence for Smart Health (KCSH), King Abdullah University of Science and Technology (KAUST), Thuwal 23955-6900, Kingdom of Saudi Arabia; Center of Excellence on Generative AI, King Abdullah University of Science and Technology (KAUST), Thuwal 23955-6900, Kingdom of Saudi Arabia

**Keywords:** Short tandem repeats (STRs), Genome annotation, Population variation, Cancer genomics

## Abstract

Tandem repeats (TRs) are highly polymorphic genomic elements, associated with diverse molecular traits and implicated in numerous human diseases. However, large-scale analysis of TRs has been limited by computational challenges, including motif recognition, detection in complex regions, and excessive computational cost. Here we present FastSTR, a computationally efficient tool for precise detection and characterization of TRs. FastSTR integrates a context-aware N-gram motif model with a segmented global alignment algorithm to enable accurate motif identification and boundary definition, even for repeat units up to 8 bp. Across 13 species, FastSTR achieved >90% recall and 99% precision, running several times faster than existing methods white outperforming them in both sensitivity and accuracy. Applied to the human genome, FastSTR uncovered previously unannotated HSATII elements, resolved population-specific TR demonstrate, and identified recurrent STR alterations in lung cancer. These results demonstrate FastSTR as a versatile framework for TR annotation and discovery, advancing studies of genome evolution, genetic diversity, and disease.

## Introduction

Short tandem repeats (STRs), also known as microsatellites, are tandem repeat sequences with repeat units(motifs) of 1-6 base pairs^1^, comprising approximately 3% of the human genome^2^. STRs exhibit high polymorphism, characterized by significant variations in length, sequence composition, and copy number^3,4^. Their mutation rates are estimated to be 10 to 100,000 times higher than those of other genomic regions, reflecting high instability^5,6^. These characteristics make STRs highly valuable information to be considered in forensic science, paternity testing, population genetics, and genetic disease diagnosis^7,8,9^. In forensic investigations, comparing STR profiles from crime scene evidence to those of suspects can establish identity and provide compelling legal evidence^10^. Due to their Mendelian inheritance, comparing the STR profiles of a child and potential parents can confirm parentage^11^. Research has shown that approximately 60 STR loci are associated with various Mendelian disorders^12^, including ataxias^13^, amyotrophic lateral sclerosis^14^, Huntington’s disease^15^, fragile X syndrome^16^, and numerous neurological conditions^17,18,19^. Furthermore, STR expansions have been linked to at least seven types of cancer, including lung and kidney cancers^20,21^.

Since the late 1970s, techniques such as Southern blotting^22^, Sanger sequencing^23,24^, and capillary electrophoresis^10,25^ have been used to identify STRs loci. While these methods are generally accurate, they are labor-intensive and time-consuming, limiting their scalability for whole-genome analysis and large cohort studies^26,27^. The rapid advancement of high-throughput sequencing technologies has laid the foundation for genome-wide STRs detection^28,29,30^. Various methods using sequencing data or assembled genomic sequences have been developed for this purpose, which can be categorized into database-based and de novo detection approaches^12,31^. Database-based methods typically identify STRs that share significant similarity with known sequences, and thus fail to detect unannotated or novel STRs^32,33^. In contrast, de novo approaches do not rely on pre-defined repeat sequence libraries, overcoming the limitations of reference dependence and enabling broader application in practical detection tasks^12,31^. However, existing de novo methods still suffer from low efficiency, and unsatisfactory detection sensitivity. and accuracy.

De novo STR detection methods typically involve two main steps: using statistical strategies or heuristic algorithms to search for potential repeat sequences, followed by sequence alignment to generate repeat structure statistics (e.g., match percentage, indel percentage) and filtering to identify STRs with statistically significant repeat structures^34,35^. Based on these steps, de novo detection methods can be further classified into the following three categories: 1) Dictionary-based methods: These methods determine the boundaries and motif of STRs based on the distribution and relative position of seeds. Notable tools include Tandem Repeat Finder (TRF)^34^, which is widely used for searching STRs in genomes and raw sequencing reads; T-REKS^36^, which employs a K-means clustering algorithm based on seed distribution; ExpansionHunter Denovo (EHdn)^37^, which identifies STR structures and locations by scanning seed distributions; TRASH^38^, which detects motifs and their higher-order structures through seed sampling and mapping. Other examples include RepeatMasker^39^, PacmonSTR^40^, and TRstalker^41^. 2) Unique k-mer counting methods: These methods utilize sliding windows and heuristic k-mer searches to directly determine motif and STR boundary. Mreps^35^ identifies the maximum tandem repeat regions by iteratively scanning sequences and extending k-mers at both ends. STRling^42^ employs k-mer counting to identify reads enriched with short tandem repeats (STRs). By analyzing the frequency and patterns of these k-mers, it determines representative repeat units and, in conjunction with alignment data, detects STR expansions at both known and novel loci. Additional tools include HipSTR^43^, GMATo^44^, and TRDistiller ^45^. 3) Machine learning-based or deep learning-based methods. These approaches train predictive models to identify STRs by extracting sequence features (e.g., motif periodicity, entropy) or directly learning latent representations from raw sequencing data. The trained models then detect STRs through pattern recognition or end-to-end prediction. RExPRT^46^ uses an ensemble method comprising support vector machines (SVM) and extreme gradient boosting (XGBoost) to predict pathogenic STRs; DeepRepeat^26^ is a CNN-based program that identifies STRs directly from nanopore electrical signals; DeepTRs^47^ employs multimodal transformation and CNNs to detect STR variants from raw nanopore sequencing reads. Other examples include WarpSTR^48^, DeepSymmetry^49^, and NanoSTR^50^.

Although the aforementioned detection methods generally fulfill the requirements for STR detection, significant challenges remain in handling diverse STR detection contexts, such as distinguishing STRs in high-variation regions, and detecting novel STRs with varying repeat units^12,51^. Moreover, detection strategies often struggle to balance sensitivity, precision, and computational cost. First, most tools have limitations in terms of recognition length and precision, preventing comprehensive detection of various STRs. lobSTR^52^ exhibits higher error rates when processing certain dinucleotide repeats, such as AC/TG repeats, which adversely affects its accuracy at these loci. RepeatSeq^53^ performs suboptimally in detecting longer STRs or expansion events, thereby limiting its effectiveness in identifying certain disease-associated STRs; mreps^35^ cannot identify STRs containing insertions/deletions (indels); TRF^34^ lacks an accurate statistical significance measurement, causing interruptions when identifying STRs with high local variation—an issue also presents in Straglr^54^. Machine learning and deep learning-based methods, while more tolerant to noise and sequencing errors, often produce a large number of novel STR predictions, many of which may be false positives^46, 47^. Secondly, many tools have overly complex parameter settings and their performances are highly dependent on parameter choices, making them less adaptable to diverse detection scenarios compared to methods that allow flexible parameter settings and have lower dependence on hyperparameter tuning. For example, T-REKS’s^36^ detection accuracy is highly sensitive to the selection of parameter K, while TRASH^38^ may require adjustment of up to 10 parameter options for varying precision levels. Last but not least, most current tools require substantial computing resources and many lack support for parallel execution or multi-threaded processing. For instance, traditional motif recognition and sequence alignment algorithms often exhibit high time complexity, resulting in excessive runtime when processing long sequences^26,34, 36^. These challenges highlight the need for further improvement in STR detection methods, particularly the unmet need for time- and computation-efficient algorithms that can keep pace with the rapid expansion of genomic data.

Here, we present FastSTR, an STR detection tool designed for efficient motif recognition and sequence alignment to rapidly identify STRs in genomic sequences while maintaining high precision and completeness. FastSTR follows a de novo assembly strategy and includes a density clustering-based fuzzy repeat region identification module that can filter out non-STR regions while retaining segments with significant local variations. It also features a context-aware motif recognition module and a high-speed segmented global sequence alignment module. To enhance detection speed, FastSTR partitions the genomic sequence into overlapping subsequences for parallel processing. Compared to the existing STR detection tools, FastSTR is not only highly efficient and easy to configure, but also does not require predefined motif libraries or rigid repeat-length thresholds, making it adaptable to a variety of STR detection scenarios.

## Results

### Overview of FastSTR

While the continuously decreasing cost of high-throughput sequencing enables the generation of massive genomic data, the rapid detection of genetic variants—including short tandem repeats (STRs)—from large-scale datasets remains an unmet need^12,55^. Here, we designed FastSTR to accurately identify STRs from genomic sequences in a time- and computation-efficient manner.

FastSTR operates in three main stages: (1) segmenting sequences from the input file (e.g., FASTA file), (2) detecting STRs within the segments using parallelized detection modules, and (3) merging the results from multiple segments to produce the final identification output (Figure 1a). At each stage, FastSTR employs various strategies to ensure efficient STR detection. Initially, it segments the input genomic sequences into fixed-length fragments with overlapping ends and further processes these fragments in parallel within the detection module (Figure 1b). Within each segment, FastSTR first identifies anchors—pairs of identical bases separated by a predefined distance—and uses their distribution to detect fuzzy repeat intervals via density clustering (Figure 1c). Next, FastSTR calculates the density of leading bases (the earlier bases in anchors) within each fuzzy repeat interval, filtering out a substantial number of intervals that lack significant tandem repeat structure. The remaining intervals are classified into trivial repeat regions containing a single STR or obfuscation repeat regions containing multiple STRs. FastSTR then applies a motif detection algorithm based on the N-Gram model and Markov chains to identify representative motifs for each interval (Figure 1d), mapping them back to their respective intervals. Intervals with low mapping quality are further filtered out. For complex intervals, a density-based refinement method (“Concentration Tests”) is used to delineate the approximate positions of genuine STRs. For each filtered fuzzy repeat interval and its corresponding putative STR, FastSTR refines its boundaries using probe alignment (probes formed by STRs of specific lengths) and sequence merging strategies, preparing these candidate STR sequences for a final alignment step to accurately determine their exact boundaries and obtain detailed alignment information. For ultra-long STRs, FastSTR employs a segmented global sequence alignment algorithm to significantly reduce computational time (Figure 1e).

**Figure 1.**
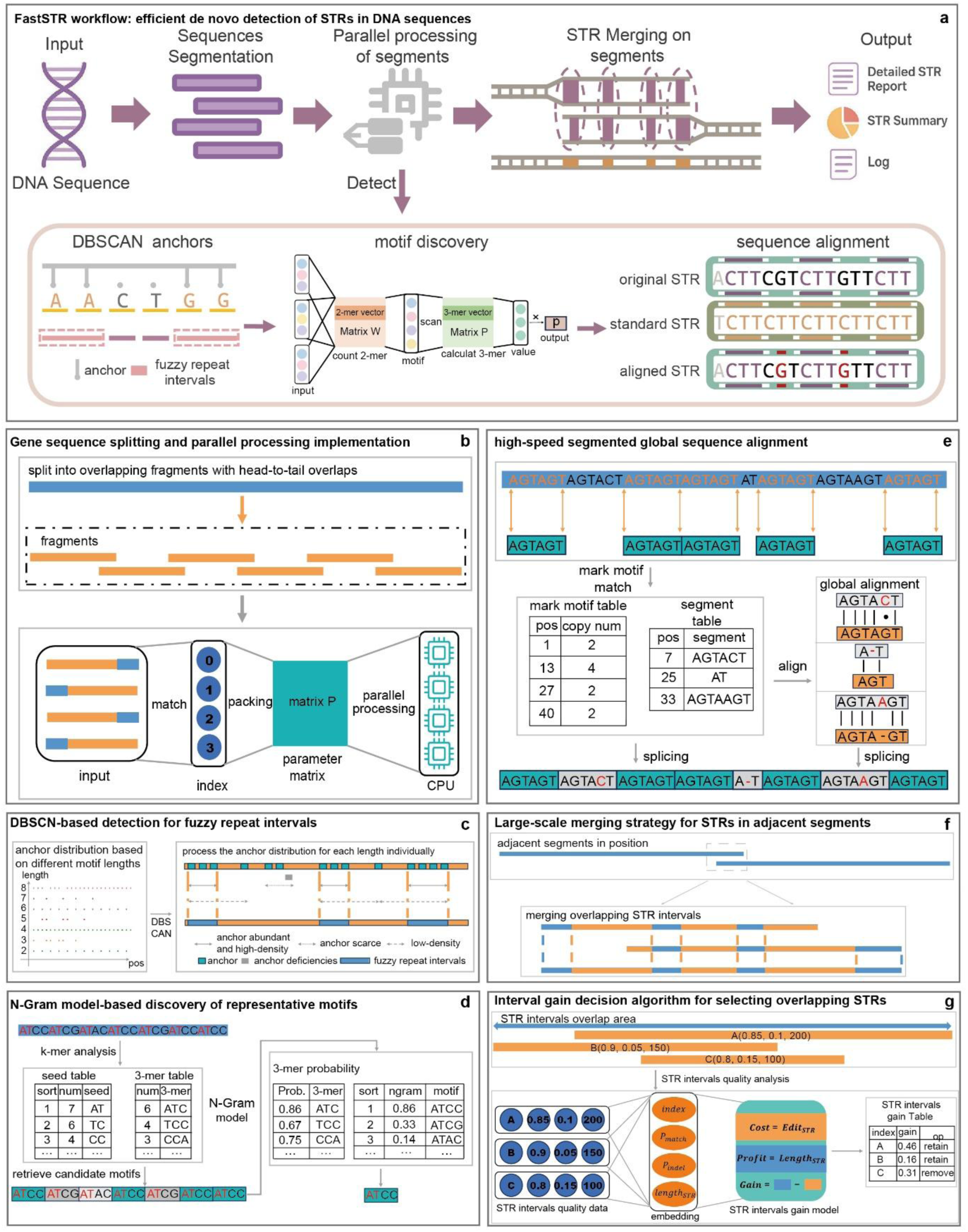
A schematic of FastSTR. **a**, FastSTR slices the input genome into segments and performs parallelized STR detection to greatly accelerate runtime. **b**, The genome is partitioned into overlapping sequence fragments. **c**, Fuzzy repeat regions are identified using density-based clustering. **d**, The most probable motif for each region is inferred using an N-gram model combined with a Markov chain. **e**, Ultra-long STRs are efficiently aligned using a segmented global alignment algorithm. **f**, STRs detected in adjacent fragments are merged if they overlap. **g**, An interval gain algorithm resolves overlapping STRs by selecting the most informative calls.

In the final step, FastSTR resolves overlapping STRs across different fragments using a “large-scale merging strategy” combined with an overlap selection strategy based on an interval gain decision algorithm (Figure 1f,g). During this process, FastSTR efficiently constructs consensus sequences while handling ultra-long STRs spanning multiple fragments using parallel alignment methods, further enhancing processing speed.

### Construction of robust ground truth

To evaluate the accuracy of FastSTR, we constructed a dataset by extracting STR loci from T2T-CHM13v2.0 assembly^28^ based on TRF-derived annotations^34^. We further examined the quality of STRs in this dataset to ensure a robust ground truth. We defined two metrics: the match percentage (*P*_*Match*_) and the indel percentage (*P*_*Indel*_) (see Methods). The joint distribution of *P*_*Match*_ and P_Indel showed that 99.69% of STRs achieved *P*_*Match*_>0.80 and *P*_*Indel*_<0.15, whereas only 0.22% fell below the match threshold and 0.09% exceeded the indel threshold. As P_Indel increased or P_Match decreased, the quality distribution derived under TRF’s default settings converged sharply around the cutoffs *P*_*Match*_ > 0.80 and *P*_*Indel*_ < 0.15, with minimal variance. This indicates that these thresholds effectively discriminate high-quality STRs from lower-quality or spurious calls.

We further performed the same analysis separately for STRs of each motif length (Supplementary Fig. 2,3), confirming the thresholds’ generalizability across diverse genomic contexts. Building on these findings, we introduced a composite quality index

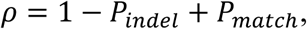

to summarize individual STR quality. Analysis of the ρ distributions for the full dataset and each motif category (Fig. 1; Supplementary Fig. 4) revealed the near absence of STRs simultaneously exhibiting high indel and substitution rates (1 − 0.15 + 0.80 = 1.65 < 1.75 ≈ min(1 − *P*_*indel*_ + *P*_*match*_)). Moreover, distinct motif lengths displayed statistically significant shifts in *ρ*, reflecting inherent sequence-specific quality profiles.

Finally, to assess the impact of alignment tolerance and score thresholds on quality metrics, we reconstructed the test dataset under relaxed alignment parameters and lower score cutoffs and compared the resulting *P*_*Match*_, *P*_*Indel*_, and *ρ* distributions to those obtained with default TRF settings. The distributions remained highly concordant, indicating that regardless of parameter variation, STR quality profiles are predominantly governed by TRF’s underlying scoring model (*P*_*I*_ = 0.10, *P*_*M*_ = 0.80), leading to minimal shifts in quality distributions.

Using the same strategy, we constructed STR ground truth sets for additional 12 species, including human, wheat and drosophila (full list in Table 1). Except for Drosophila melanogaster (xx Gb), rice (< 0.5 Gb) and wheat (13.75 Gb), most genomes were approximately 2 Gb in size. However, the total number of STRs varied substantially across species, ranging from 15,276 to 946,316. Notably, maize had the lowest number of STRs among the 13 species, whereas wheat - despite its larger genome - had an STR count comparable to that of human. In addition, the mouse genome contained nearly one million STRs—significantly more than human and other mammals. These interspecies differences in STR abundance likely reflect distinct genome architectures and evolutionary mechanisms underlying STR formation and maintenance (Supplementary notes).

### Accuracy of FastSTR

To comprehensively evaluate FastSTR’s detection performance, we ran FastSTR along with three other widely used methods, including mreps^35^, T-reks^36^, and TRASH^38^, on thirteen reference genomes and benchmarked their outputs against TRF-derived ground truth (see Methods). Across tools, FastSTR’s STR counts most closely approximated the ground truth, with the ratio of detected to ground-truth STRs remaining around 1.3 regardless of genome size. In comparison, mreps exhibited a ratio between 6.4 and 45.7, and T-reks between 58.1 and 383.5, indicating that FastSTR delivers a more stable and consistent STR detection profile. TRASH yielded a comparatively low STR count because it focuses on higher-order repeats and cannot characterize STRs with repeat units shorter than 7 bp^38^.

We next evaluated the recall and precision of STR detection by FastSTR and benchmarked its performance across thirteen diverse genomes. Overall, FastSTR outperformed all other tools in both recall and precision, resulting in substantially higher F1-scores across species (Figure 2d). FastSTR achieved consistently high recall rates across most genomes. Specifically, recall in animal genomes was typically above 94%, while it maintained around 90% in plant and insect genomes, demonstrating the robustness and broad applicability of FastSTR. While the highest recall was observed in the Old World monkey genome (97.03%), FastSTR also achieved high recall for human (95.3%) and mouse (Algerian mouse (96.72%) and house mouse (96.58%)), the most extensively studied model organism. On the other hand, The lowest recall was found in maize (79.57%), where missed STRs were predominantly (95.24%) those with motif lengths ≥3 bp. This is likely attributable to long-motif STRs in maize, which challenged the flanking-sequence density threshold used by FastSTR (see Methods). In terms of precision, FastSTR demonstrated exceptional performance, exceeding 99% across all genomes.

**Figure 2.**
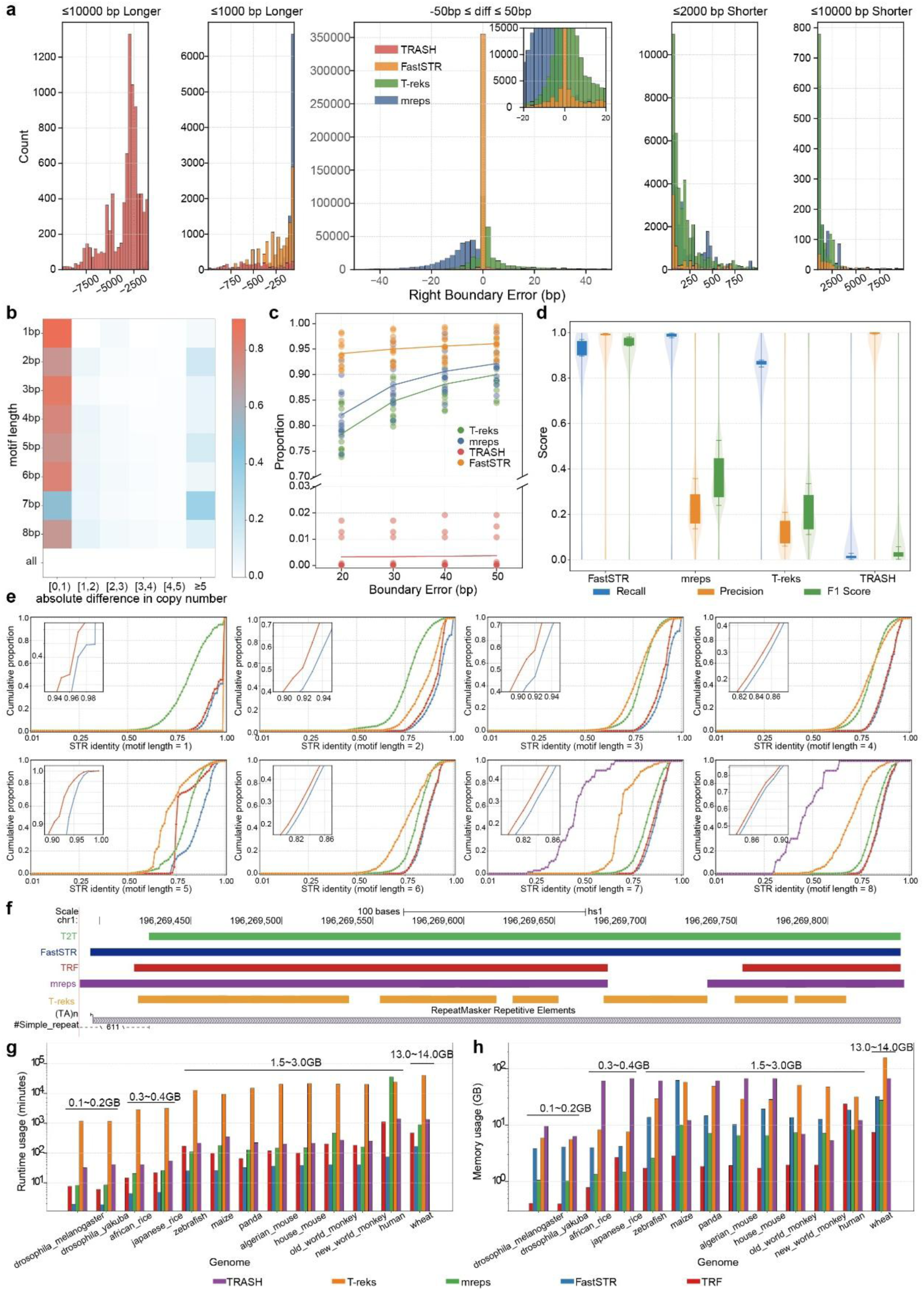
Evalutation of FastSTR. **a**, Distribution of left boundary deviation errors stratified by STR length in the human genome. **b**, Heatmap of the absolute differences in STR copy numbers between loci identified by FastSTR and Tandem Repeats Finder across the human genome, restricted to STRs sharing identical motif sequences. **c**, Proportion of detected STR boundaries falling within various boundary deviation thresholds. **d**, Performance comparison (recall, precision, and F1-score) of FastSTR, mreps, T-reks, and TRASH on 13 representative genomes. **e**, Normalized edit distance between STRs identified by all tools (including Tandem Repeats Finder) and their corresponding reference sequences. Reference sequences were constructed by perfect tandem concatenation of representative motifs without errors. STRs are grouped and summarized according to motif length. **f**, FastSTR accurately identifies STRs that are fragmented or truncated in Tandem Repeats Finder and other tools. These STRs are benchmarked against the T2T STR reference annotation set. **g**, Runtime of each tool on 13 representative genomes. **h**, Peak memory usage of each tool on 13 representative genomes.

The slightly lower recall compared to precision reflects a deliberate design trade-off that prioritizes specificity to ensure the reliability of identified STRs. By contrast, mreps exhibited high recall, typically above 98%. However, its precision was substantially lower, ranging from 13.64% in Drosophila melanogaster to a maximum of 35.67% in the Algerian mouse. T-reks showed a similar pattern: most genomes had recall above 86% but precision was mostly under 10%, peaking at just 20.84% in zebrafish. These results suggest that while mreps and T-reks capture a substantial fraction of true STRs, they produce a large number of false positives due to relaxed repeat criteria and limited discriminative power^56^.

Conversely, TRASH showed extremely high precision—exceeding 99% across all genomes. However, its recall was extremely low, generally below 1%, with the highest value observed in zebrafish (10.36%). This performance is a result of TRASH’s focus on high-order repeats, typically with motif lengths of 7–8 bp^38^. Overall, FastSTR consistently achieved much higher F1-scores compared to other methods across 13 species, highlighting FastSTR’s balanced performance in achieving both accuracy and robustness in STR detection.

We next assessed the boundary of the identified STR. For each ground-truth STR locus, we collected all overlapping STR calls from each method and calculate their shift to the ground truth locus. FastSTR consistently outperformed the other tools at all tested error thresholds (Fig 2c). Under the strictest 20 bp cutoff, over 90% of loci in nearly every genome were correctly localized, with Drosophila melanogaster even reaching >99%. When allowing up to 50 bp margin of error, most genomes exceeded 95% coverage. We further calculated the average coverage across all species under each error threshold. As the threshold increased, FastSTR’s coverage curve declined more gradually, indicating that most of its boundary errors fall within 20 bp. This superior performance arises from our boundary-refinement strategy (see Methods), which trims ambiguous flanking sequences and then applies a fine-grained alignment to precisely delineate repeat boundaries. We note that at the same error threshold, humans and zebrafish exhibited slightly lower coverage (92.8% at the 50 bp threshold; Supplementary Table S3). This is likely due to clusters of adjacent STRs with identical repeat motifs in these genomes, which prevent FastSTR from precisely distinguishing their boundaries. In comparison, mreps achieved approximately 90% coverage at the 50 bp threshold, ranking second, while T-REKS reached nearly 90% only in certain species (e.g., African rice, fruit fly, Old World monkey, and panda; Supplementary Table S3). TRASH performed the worst, with boundary coverage typically below 1%, as it focuses on identifying higher-order, long tandem repeats, whose predicted coordinates often deviate well beyond 50 bp from the ground truth^38^.

Additionally, we examined the distribution of left- and right-end boundary errors for each method in every genome. The patterns were similar and symmetric for left and right ends (Figure 2a and supplementary Figure 10). FastSTR errors were peaked at 0 bp, demonstrating highly accurate localization consistent with ground-truth boundaries. mreps showed a leftward bias (≤40 bp), reflecting its relaxed criteria for short STRs^35^, while T-reks exhibited a rightward bias (≤40 bp), indicating its tendency to fragment long STRs into shorter units^36^. TRASH errors were mainly distributed between 1,000 and 10,000 bp, due to its focus on detecting high order, very long tandem repeats^38^.

We further evaluated the concordance of copy number estimates between FastSTR and the reference. In most species, more than 80% of STRs exhibited copy number differences of no more than one unit (Fig. 2b, Supplementary Table S8). STRs with deviations greater than five repeat units accounted for less than 10% in the majority of species (Supplementary Table S8). Notably, the human genome displayed a pronounced anomaly in STRs with 7-bp motifs, with approximately 36% showing deviations greater than five repeat units. These STRs were predominantly located in the ATATATA tandem repeat arrays of the Y chromosome centromere, a structurally complex region where true STR boundaries are difficult to define ^30^. After excluding the Y chromosome, 89.5% of 7-bp motifs showed a copy number difference within 5, compared to 63.3% previously. Overall, these results indicat that FastSTR achieves accurate copy number estimation across diverse genomic contexts and thus provides a reliable foundation for exploring the biological roles of STR copy number variation.

To assess the accuracy of motif identification, we measured the edit distance between STR sequences called by each method and their corresponding ground-truth STR sequences. For each motif length (1 bp to 8 bp), we constructed a test set of 5,000 STRs per method from the human genome and further calculated the sequence identity for each identified STR.TRASH was included in the comparison only at motif lengths of 7 bp and 8 bp^38^, as it exclusively detects STRs with those motif lengths. FastSTR consistently achieved the highest sequence similarity among all methods across all motif lengths, with the most pronounced advantage at 2 bp and 3 bp motifs (Fig. 2e). This improvement may stem from the N-Gram model’s tendency to prioritize the most frequent candidate motif in short-repeat contexts, thereby minimizing edit-distance inflation due to base mismatches.

In addition to direct comparisons with TRF, we next benchmarked FastSTR-identified STRs against the T2T reference STR annotation, which represents a refined version of the TRF results with much more comprehensive manual curation. In multiple representative cases, FastSTR identified continuous repeats as a single, complete STR, in agreement with the reference annotation, whereas commonly used tools such as TRF, mreps and T-REKS tended to fragment these regions into multiple segments (Fig. 2f). In total, we identified 271 such instances genome-wide (Supplementary Table S7), highlighting the superior ability of FastSTR to preserve the integrity of STRs in complex repeat regions.

### Efficiency of FastSTR

Besides accuracy, we also assessed efficiency of FastSTR by comparing the runtime and memory usage of each tool across representative genomes. For tools lacking parallelization (TRF, mreps, and T-reks), we recorded their total runtime and peak memory usage on each genome. For parallelizable tools such as TRASH and FastSTR, we executed them on a 72-core CPU cluster and monitored memory consumption using smem^56^ (see Methods for details).

FastSTR’s execution time is markedly lower than that of other methods and increases only modestly with genome size (Fig 2g). For example, on the Drosophila genome (0.14 Gb), FastSTR completes over three times faster than TRF (2 minutes vs. 7 minutes). This also holds true for wheat (13.75 Gb), where FastSTR finishes in 2 h 43 minutes compared to TRF’s ∼8 h. These results underscore FastSTR’s time efficiency across small to large genomes. Moreover, FastSTR’s runtime scales smoothly with genome size, owing to its predominantly linear-time algorithms. Even when aligning ultra-long STRs, the segmented global alignment strategy substantially reduces alignment overhead, preventing large runtime fluctuations(Supplementary Fig. 9). To further validate the efficiency of segmented global alignment on long STRs, we generated 3,000 synthetic STR sequences (ranging from 7,000 bp to 70,000 bp in length, with motifs of 2–8 bp) and compared segmented global alignment with standard local alignment. While maintaining high alignment quality (99.57% of STRs satisfied |*ΔP*_*indel*_| + |*ΔP*_*match*_| < 0.06), segmented global alignment saved an average of 58.694 seconds per sequence, with 89.60% of sequences achieving a time reduction greater than 5 seconds. These findings confirm the algorithm’s advantage for long repeats. Although FastSTR achieves the fastest speed among all methods, its memory consumption remains at a moderate level, even showing lower memory usage than TRF on the human genome, while TRF is overall the most memory-efficient method (Fig 2h).

### A comprehensive STR catalogue generated by FastSTR

After validating FastSTR across 13 reference genomes, we further applied it to build a comprehensive STR catalogue based on the human T2T (Telomere to Telomere) genome ^58^. While STR elements identified by FastSTR showed high concordance with those (Methods) from the T2T reference annotation (604217, 95.46%) and TRF (388148, 95.03%), FastSTR detected substantial more method-specific STRs (FastSTR: 18,758, Reference: 497, TRF: 6476, all STRs filtered based on a minimum alignment score of 50), suggesting its enhanced comprehensiveness in STR identification (Fig. 3a).

**Figure 3.**
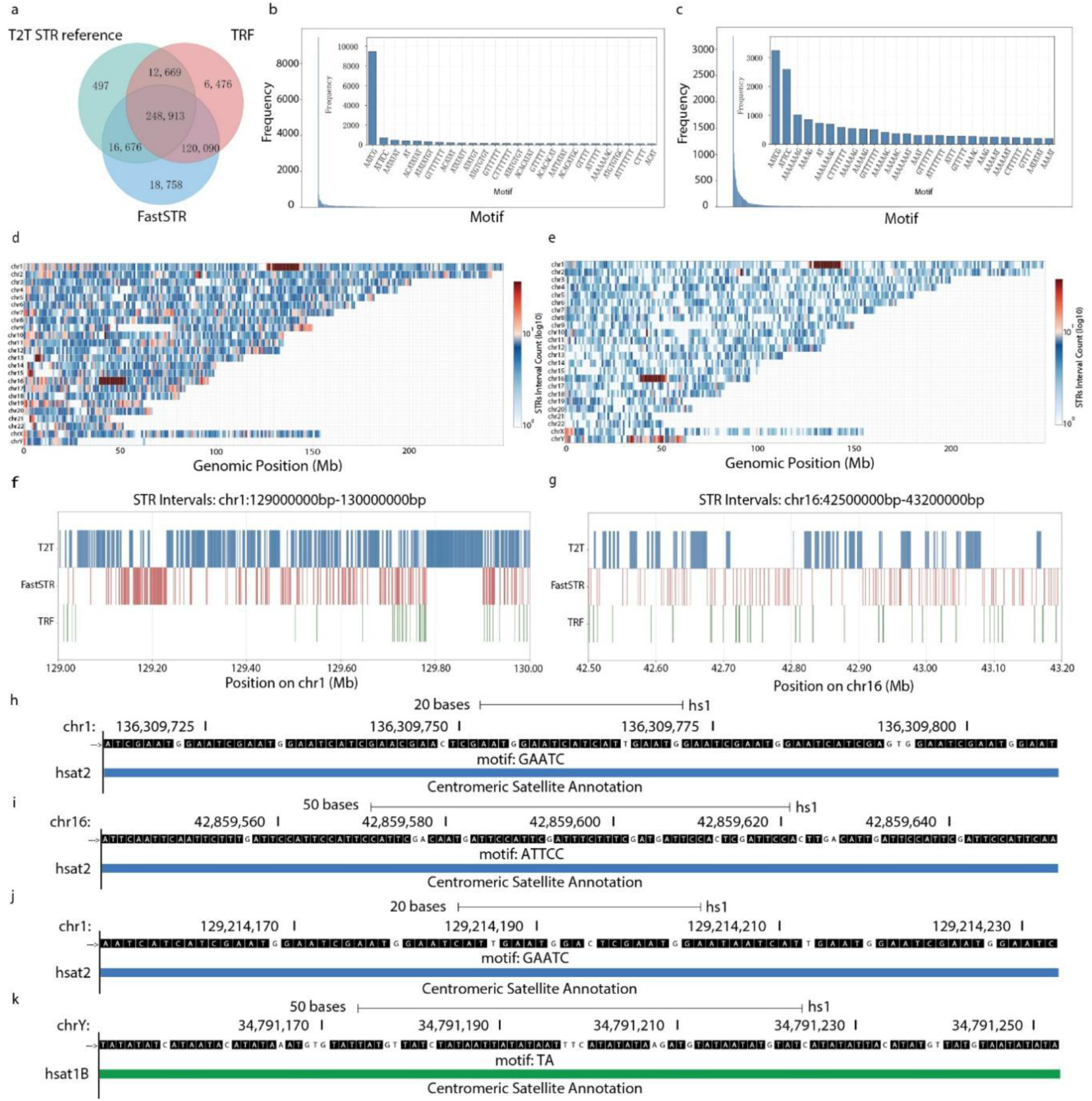
FastSTR complements existing STR reference annotations. **a**, Venn diagram showing the overlap of STR intervals detected by FastSTR, TRF, and the T2T STR reference annotation. **b-c**, Motif composition of STRs identified by FastSTR but missed by TRF; **b** shows STRs validated by the T2T reference annotation, and **c** shows novel STRs uniquely identified by FastSTR. **d-e**, Genomic distribution of TRF-missed STRs recovered by FastSTR; **d** shows novel STRs uniquely identified by FastSTR, and **e** shows those validated by the T2T annotation. **f-g**, STRs in the HSATII satellite regions on chromosomes 1 **f** and 16 **g** detected by FastSTR but absent from the T2T STR reference set. **h–k**, Representative examples of TRF-missed STRs recovered by FastSTR. **i**, **j**, STRs in the HSATII regions that complement the T2T STR reference annotation. **h**, **k**, Additional STRs recovered by FastSTR and supported by the T2T annotation.

We further dissect features of STRs detected by FastSTR but not by TRF, which were categorized into T2T-verified novel STRs (supported by the Reference) and unverified novel STRs. Verified STRs were predominantly composed of AATCG and ATTCC (Fig. 3b), corresponding to Human Satellite II (HSATII) enriched in centromeric regions (Fig. 3e)^59,60^. Unverified STRs also showed enrichment for AATCG/ATTCC, demonstrating FastSTR’s ability to recover HSATII loci that are not reported by TRF. Furthermore, FastSTR identified a novel class of motifs exhibiting structural characteristics analogous to 1–2 bp short tandem repeats (Supplementary Fig. 15), expanding STR diversity. These motifs were undetectable by TRF. To map STR distributions, we divided the T2T genome into 1 Mb bins and quantified STR density per bin (Fig. 3d, 3e). Compared with TRF, FastSTR substantially augmented HSATII coverage in the centromeres of chromosomes 1 and 16, and detected additional STRs in AT-rich HSATIB regions of chromosome Y^30^, all validated by the T2T reference annotation. Even compared with the reference, FastSTR still improved the detection of HSATII in the centromere of chromosome 1 and 16. Further analysis of HSATII localization in the centromeres of chromosomes 1 and 16 revealed that FastSTR filled extensive gaps in TRF-detected STRs the T2T HSATII annotations (Fig. 3f, 3g). Representative HSATII and HSATIB-associated STR sequences identified by FastSTR (Fig. 3h–3k), exhibited unambiguous tandem repeat structures. These results robustly demonstrate FastSTR’s capacity to enhance STR annotation in the human genome, resulting in a more comprehensive human STR catalogue.

### Identification of STR polymorphism with FastSTR

To investigate STR polymorphism, we applied FastSTR to detect STR variants in samples from the 1000 Genomes Project and Human Pangenome Project^61,62,63^, followed by population- and cohort-level analysis of genomic STR variation. The dataset comprised 82 individuals across 28 populations grouped into five super-populations: African (AFR, n=37), European (EUR, n=8), South Asian (SAS, n=6), East Asian (EAS, n=12), and Admixed American (AMR, n=21), including 35 males (42.68%) and 47 females (57.32%).

We designed a pipeline using FastSTR to detect STR variations in each individual genome assembly relative to the reference (Fig. 4a). In brief, each genome assembly was used as input to identify STR loci with FastSTR, while simultaneously being aligned to the reference genome using minimap2^64^, and variants were called with paftools^64^. The resulting variants were then intersected with the identified loci to generate genome-wide STR variation profiles using custom scripts and bedtools^65^. We further compared the FastSTR results with those derived from the T2T reference annotation, and a recently published benchmark TR catalog ^66^. STR variants were into four categories: (1) those identified by all three resources (All), (2) those identified only using the benchmark TR catalog (hg38-only), (3) those identified only using the T2T reference (T2T-only), and (4) novel variants (Novel) identified exclusively by FastSTR. It turned out that FastSTR identified substantial Novel STR variants in each individual genome (Fig. 4b).

**Figure 4.**
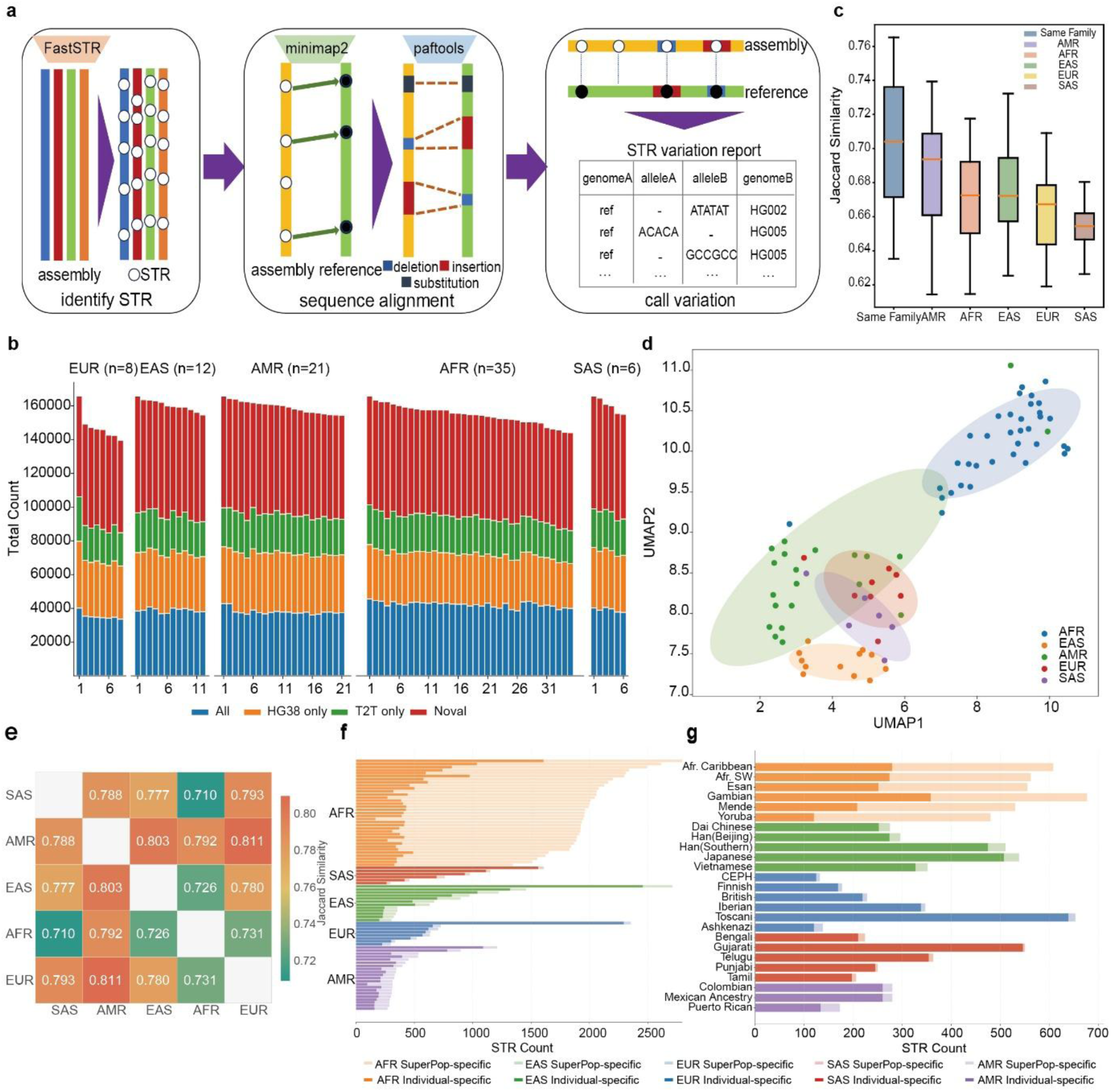
Identification of novel STR polymorphism. **a** Overview of the genomes analyzed and the pipeline used to identify STR polymorphisms. **b** Box plots showing Jaccard similarity coefficients of STR profiles within families and between unrelated families. **c** Distribution of the number of novel versus previously annotated STR variation loci. **d** UMAP projection of samples from five population groups based on STR variation. **e** Heatmap of pairwise Jaccard similarities across the five populations. **f-g** Distribution of population-specific (light color) and individual-specific (dark color) STR variation counts.

We further evaluated the reliability of FastSTR-identified variants by comparing variant profiles among individuals within the same family and across different families in each super-population. As expected, Jaccard similarity coefficients calculated from STR variant loci demonstrated significantly higher similarity among individuals within the same family versus those from different families (t=2.63–5.78, adjusted p=6.1×10⁻⁸–4.6×10⁻²). UMAP visualization of population-level STR variation (Fig. 4d) showed: (1) clear separation of AFR from other clusters, reflecting its highest genetic diversity; (2) AMR forming a continuum between African, European, and East Asian clusters, indicative of multifaceted ancestral origins (indigenous, European migrant, and African-descendant admixture); (3) proximity of EUR and SAS clusters, suggesting historical gene flow; and (4) a distinct East Asian cluster, supporting unique migratory history. Inter-population Jaccard similarity (Fig. 4e) further validated this structure: AFR showed the lowest similarity to other groups (0.71–0.73), while AMR had high similarity to EUR/EAS/AFR (0.79–0.81).^7,67,68,69^

Quantification of population-specific and individual-specific STR variants (Fig. 4f) revealed significantly more population-specific variants in AFR, implying richer variation accumulation and longer independent evolutionary history. Non-African populations showed reduced shared variation due to historical bottlenecks^63^. Individual-specific variant counts did not differ significantly across groups, being largely driven by individual mutation rates and local genomic context. Analysis of STR variants >50 bp (structural variation perspective) recapitulated identical distribution patterns (Fig. 4g). Thus, STR variation not only mirrors structural diversity but also serves as a sensitive marker for reconstructing population history and genetic architecture.

### Detection of novel STR variants in lung cancer by FastSTR

In addition to profiling STR variations across different induvial, we also aimed to apply FastTR to identify STR mutations and variants in cancer samples relative to the normal reference genome. To this end, we first performed de novo assembly on whole-genome sequencing data generated by the University of Tokyo Medical Genome Sciences Laboratory^70^ (UT-MGS; Supplementary Table 3) for 8 lung adenocarcinoma and 1 breast cancer samples. We then identified STR variants in these tumor assemblies relative to the reference genome and compared them to those from normal samples described in the previous section. In total we identified 132,808 variant sites shared across tumor and normal samples (Fig. 5a). Although the lung cancer sample size was much smaller than that of normal samples (8 vs. 82), it contained a substantial higher proportion of its specific variants (35.14% vs 24.9%), indicating an elevated frequency of STR variations in cancer (Fig. 5a). To examine this more precisely, we calculate STR density (per Mbp) and STR variant burden (per Mbp) for each genome assembly. Normal samples exhibited a mean STR density of 195.85 per Mbp compared with 162.92 per Mbp in lung cancer assemblies (Fig. 5b), likely reflecting the higher genomic coverage of normal samples from the 1000 genome project and human pangenome project. By contrast, STR variant burden per Mbp was significantly higher (p = 1.29e-11) in tumour samples (73.26) than in normal samples (40.50), confirming an elevated STR mutation load in the cancer cohort despite potential assembly-quality effects (Fig. 5c).

**Figure 5.**
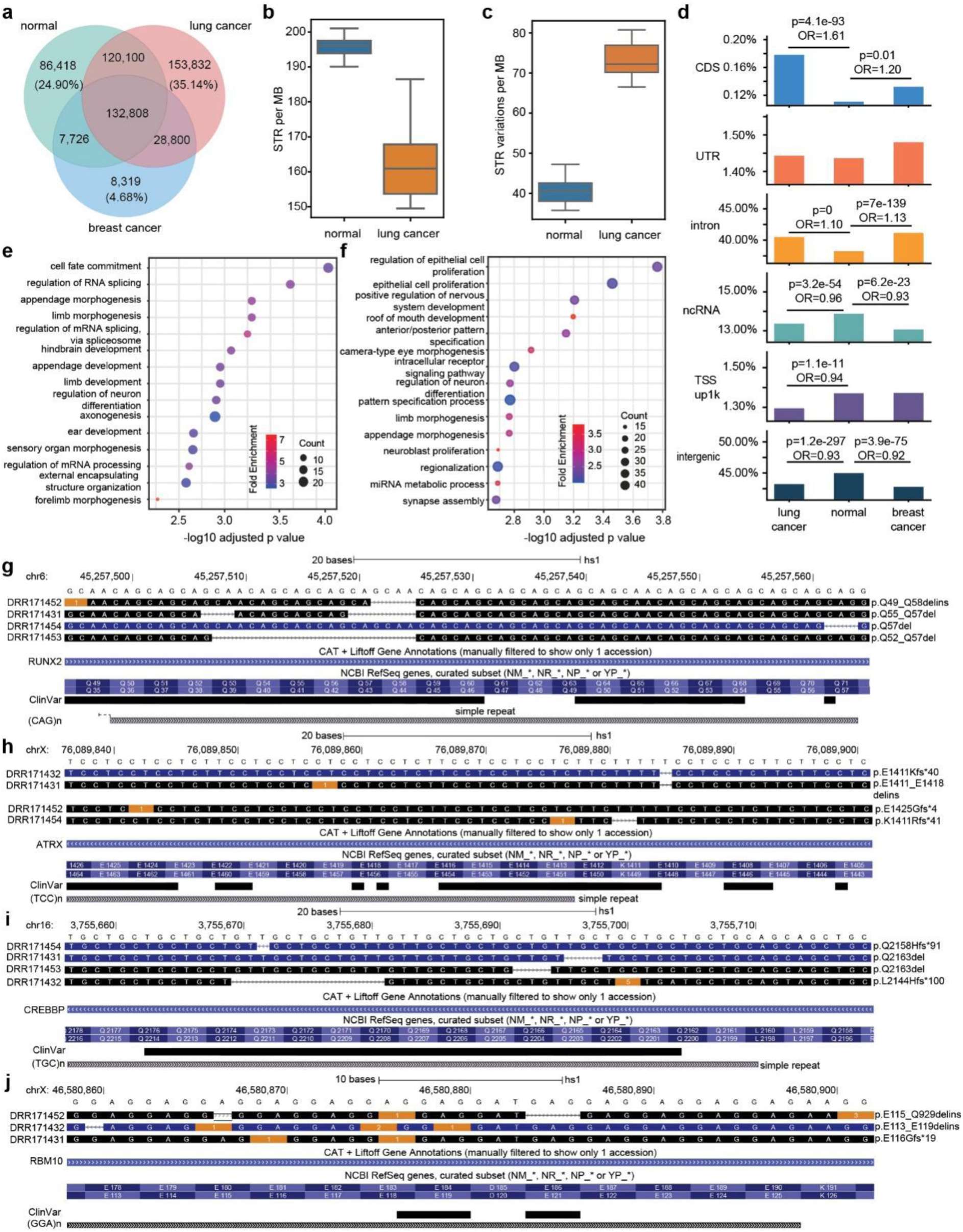
Detection of novel STR variants in lung cancer. **a**, Venn diagram showing the numbers of STR variants in normal, lung cancer, and breast cancer samples. STR counts were binned, and overlapping STRs were merged into unified intervals. **b**, Density of STRs per Mbp in lung cancer and normal samples. **c**, Density of STR variants per Mbp in lung cancer and normal samples. **d**, Proportion and statistical significance of STR variants located in different genic regions (CDS, UTR, intron, TSS_up1k, ncRNA, intergenic) across lung cancer, breast cancer, and normal samples. When an STR overlapped multiple regions, assignment followed the priority order: CDS > UTR > intron > TSS_up1k > ncRNA > intergenic. **e,f**, GO enrichment analysis (biological process, BP) of lung cancer–specific STR variants in CDS **e** and UTR **f** regions. **g–j**, Visualization of representative STR variant mechanisms in lung cancer genes using UCSC Genome Browser. Yellow blocks indicate base insertions. All genes are annotated with NCBI RefSeq transcripts, and corresponding amino acid changes are shown on the right (ANNOVAR).

We further examined the genomic distribution of these lung cancer specific STR variants by partitioning the genome into six functional classes: coding sequences (CDS), un-translated region (UTR), intron, 1000 bp upstream of transcription start site (TSS_up_1k), ncRNA and intergenic regions. Although most STR variants fall within intergenic (42.8–45.0%) and intronic (38.2–41.1%) regions, cancer samples show a marked enrichment of STR variants in functional genic regions compared with normal samples (Fig. 5d). In particular, CDS sites are the most significantly enriched for cancer STR variants (lung: p = 4.1×10⁻⁹³, OR = 1.61; breast: p = 0.01, OR = 1.20; Fisher’s exact test), and introns are also enriched (lung: p ≈ 0, OR = 1.10; breast: p = 7×10⁻¹³⁹, OR = 1.13), whereas cancer variants from ncRNA and intergenic regions are relatively depleted (for intergenic: lung p = 1.2×10⁻²⁹⁷, OR = 0.93; breast p = 3.9×10⁻⁷⁵, OR = 0.92). These patterns indicate that cancer-associated STR variation may preferentially affects functionally relevant genomic regions^21^.

We next investigated the genes harboring these STR variations and calculated their frequency across the eight tumour assemblies. In total, 466 and 1043 genes contained STR variants within CDS and UTR, respectively. Interestingly, genes with variants in CDS regions were most significantly enriched for cell fate commitment and regulation of RNA splicing, whereas those with variants in UTR were most significantly enriched for epithelial cell proliferation functions (Fig. 5e and 5f). These processes are critical for cancer development and progression.

Two representative examples of genes with variants in CDS and relevant to these functions are CREB-binding protein (CREBBP) and RNA binding motif 10 protein (RBM10). CREBBP encodes a lysine acetyltransferase that acetylates histones and plays critical roles in growth control and hemostasis by coupling chromatin remodeling with transcription factor recognition^71,72^(see GeneCards). We identified STR variants within the CDS of CREBBP in 4 out of 8 lung cancer samples, including one resulting in a frameshift near the HAT domain (Fig. 5g). This finding is consistent with previous studies showing that losss of HAT activity in CREBBP is causally linked to oncogenic programs in several tumour types ^72^. RBM10 is a validated splicing regulator recurrently mutated in lung adenocarcinoma. Its truncation or disruptive mutations have been shown to alter splicing of functionally important targets (e.g. NUMB) and to contribute to tumour phenotypes ^73,74,75^. Consistently, we identified STR variants within its CDS in 3 lung cancer samples, resulting in broader functional deficiency, including frameshift (DRR171431) and truncation/stop gain (DRR171452) and an in-frame indel (DRR171432) predicted to alter the RNA-recognition motif (Fig. 5h).

In addition to these two, there are also 463 genes with recurrent STR variants within CDS across the 8 lung cancer samples. Many of them have also been implicated in cancer-related functions. For instance, RUNX2 contains an AGC trinucleotide STR within the annotated CDS. Four tumour assemblies display copy-number loss of this STR, resulting in deletion at its N-terminal QA transactivation region. Previous functional studies have shown that QA-repeat length/structure modulates RUNX2 transactivation activity and downstream transcriptional programs, and perturbation of this domain can alter RUNX2’s oncogenic properties ^76,77^. ATRX is a chromatin-remodelling factor whose truncating mutations have been repeatedly linked to loss of function, telomere maintenance defects and the alternative lengthening of telomeres (ALT) phenotype in multiple tumour types ^78,79,80^. Indeed, we found that 4 tumor samples loss copies of a CCT STR within its coding sequence, resulting in recurrent frameshift in 3 samples.

In summary, applying FastSTR to de novo tumour assemblies reveals a substantial number of novel STR variants in lung cancer, enriched in coding and regulatory regions associated with cancer-related functions as well as transcriptional and splicing regulation (Fig. 5a–j). Several of these STR events lead to protein-altering consequences (in-frame delins, frameshifts and premature stops) in recurrently affected cancer genes, such as RUNX2, ATRX, CREBBP, RBM10 These findings suggest that STR-mediated coding perturbations represent an important contributor to lung cancer genomics and further demonstrate the potential utility of FastSTR in cancer genomics research.

## Discussion

Here we present FastSTR, a next-generation STR detection tool that achieves high efficiency and accuracy compared with existing commonly used STR detection methods. Compared with commonly used STR detection methods, FastSTR consistently achieves higher accuracy and greater time efficiency, while offering enhanced interpretability of motifs of varying lengths. These advances are enabled by two innovations: a context-aware N-gram–based motif recognition model and a high-speed segmented global alignment algorithm for ultra-long STRs. The N-gram model effectively captures both positional and sequential information of STRs, allowing linear-time identification of the most probable motifs. The segmented alignment strategy further accelerates computation by decomposing long STRs into shorter, informative fragments, markedly reducing runtime without loss of accuracy. In addition, FastSTR introduces a principled strategy for resolving overlapping STRs based on an interval-gain decision framework, thereby avoiding subjective biases and retaining biologically meaningful STRs within the same genomic interval. To address the challenge of complex centromeric satellite regions, we introduce the concept of confounding repeat regions and a novel partitioning approach (“Concentration Tests”) that enables their systematic delineation. Applied to the human genome, this strategy revealed numerous previously unannotated HSATII (Human Satellite II) elements absent from current T2T references, underscoring the importance of STR-centric approaches in refining genome annotations.

Beyond annotation, we developed a lightweight de novo STR variant detection pipeline integrating FastSTR with minimap2 and paftools, enabling sensitive detection of assembly-specific STR variation. Applied to population-level datasets, this pipeline uncovered STR variants that highlight population differentiation and evolutionary divergence, demonstrating the potential of STRs as informative markers of genetic and evolutionary processes. In cancer genomics, we applied our pipeline to lung cancer samples and identified recurrent STR variants in key cancer-associated genes. These findings reveal that STR variation represents an important mutational class in tumor evolution, opening avenues for future exploration of STR-driven mechanisms in oncogenesis.

We conducted a comprehensive evaluation of FastSTR across the genomes of 13 species. Interestingly, we observed substantial variation in STR abundance across species. For example, maize contained only ∼50,000 STRs, a number strikingly lower than that of species with similarly sized genomes, whereas wheat harbored a genome several times larger than human yet with a comparable number of STRs. In contrast, mouse displayed nearly one million STRs, greatly exceeding that of humans and other mammals. These patterns suggest fundamental biological differences in STR density across kingdoms. A plausible explanation is that plant genomes may rely more heavily on other types of repetitive sequences (e.g., transposable elements) for genome expansion, whereas rodent genomes may have undergone STR expansion through lineage-specific mechanisms. Furthermore, recall rates of the FastSTR stabilized around 95% for animal genomes and ∼90% for plants and insects, suggesting species-specific distributions of STR motif lengths and quality profiles^81,82^. Notably, STRs with motifs of 7–8 bp differed markedly from those with 1–6 bp motifs in both quality distribution and recognition accuracy. This observation may reflect the conventional definition of short tandem repeats (STRs), which are typically restricted to repeat units of 1–6 bp. Such a boundary is not arbitrary but rather grounded in their distinct mutational mechanisms and evolutionary dynamics, which confer biological properties that differ markedly from those of longer repeat units^81,83^.

We observed that FastSTR was able to identify a large number of previously unannotated STRs, the majority of which were HSATII (Human Satellite II) repeats located at the centromeric regions of chromosomes 1 and 16. In addition, FastSTR detected a specific class of STRs with motif structures composed of alternating 1 bp and 2 bp repeat units, which are often missed by tools such as TRF.

Interestingly, cancer-associated STR variants were predominantly deletions rather than insertions. We hypothesize that this reflects a genuine biological signal rather than a technical artifact, consistent with prior studies suggesting that cancers frequently accumulate deletion-type STR mutations as part of their mutational processes ^84^. It is important to note that our STR de novo variant detection may be subject to multiple sources of bias. First, detection of variants by minimap2 and paftools depends on alignment parameters; second, the coordinate projection performed by paftools can introduce systematic error; third, the assembly quality of lung cancer samples may influence the false discovery rate of variants; and finally, since STRs frequently overlap, we chose to merge overlapping STRs when quantifying variants, a strategy that is widely adopted in current practice.

Despite addressing key challenges of accuracy and speed in STR detection across multiple species, several limitations of FastSTR remain. First, the memory footprint can still be substantial when analyzing highly complex genomes. This is likely attributable to the segmented global alignment algorithm used for mapping long STRs, which requires significant heap memory. Future versions will aim to optimize this strategy to achieve low-memory, time-efficient, and high-accuracy alignment of ultra-long STRs. Second, FastSTR faces difficulties in fully resolving highly complex centromeric satellite regions, where extended microsatellite arrays are often only partially detected. This limitation reflects a broader challenge faced by all STR detection tools, as these regions frequently contain higher-order satellite repeats whose fundamental repeat units are themselves tandem repeats, complicating the precise definition of STR boundaries. To address this, we plan to refine the proposed segmentation method and explore satellite-specific assembly algorithms, with the goal of enabling more comprehensive characterization of these regions in future releases.

In summary, we present FastSTR as a versatile and high-performance tool for the detection and analysis of STR across diverse species and genomic contexts. FastSTR not only achieves high accuracy and efficiency in motif recognition, boundary delineation, and copy number estimation, but also enables scalable population- and disease-level analyses, revealing biologically meaningful STR variation and patterns that have been previously underexplored. By combining methodological innovations such as the N-gram–based motif model, segmented global alignment, and interval-gain decision framework, FastSTR provides a robust framework for both comprehensive STR annotation and de novo variant discovery. We anticipate that FastSTR will facilitate future investigations into the roles of STRs in genome evolution, population differentiation, and disease, and that its principles may be broadly applicable to other species and complex genomic regions, thereby expanding the landscape of repeat-based genomic research.

## Online content

## Publisher’s note

Springer Nature remains neutral with regard to jurisdictional claims in published maps and institutional affiliations.

Springer Nature or its licensor (e.g. a society or other partner) holds exclusive rights to this article under a publishing agreement with the author(s) or other rightsholder(s); author self-archiving of the accepted manuscript version of this article is solely governed by the terms of such publishing agreement and applicable law.

© The Author(s), under exclusive licence to Springer Nature America, Inc. 2024

## Methods

### Samples and preprocessing

To comprehensively evaluate the efficiency and accuracy of FastSTR in detecting short tandem repeats (STRs), we selected 13 representative genomes across diverse taxa: human (T2T-CHM13v2.0), zebrafish (GRCz11), Drosophila melanogaster (Release 6 plus ISO1 MT), Drosophila yakuba (Prin_Dyak_Tai18E2_2.1), maize (Zm-B73-REFERENCE-NAM-5.0), Old World monkey (ASM2454274v1), New World monkey (mCalJa1.2.pat.X), house mouse (GRCm39), Algerian mouse (SPRET_EiJ_v3), giant panda (ASM200744v3), Japanese rice (AGIS1.0), African rice (OglaRS2), and wheat (IWGSC CS RefSeq v2.1). All genome assemblies were obtained from the NCBI database (Supplementary Table S1).

We employed Tandem Repeats Finder (TRF)^34^ to identify all tandem repeat sequences within each genome, filtering for STRs with repeat unit lengths of 8 base pairs or fewer. To assess FastSTR’s capability in detecting overlapping STRs, we retained overlapping regions identified by TRF. TRF was executed with default parameters, except for alignment settings, which were adjusted from “2 7 7” to “2 5 7” to lower the detection threshold and capture a broader range of STRs.

We downloaded tandem repeat annotations for the human T2T-CHM13v2.0 genome from the UCSC Genome Browser^58^ (as defined by RepeatMasker) and filtered for STRs with repeat unit lengths of ≤8 bp to construct a comprehensive STR annotation set for T2T. In addition, we obtained a tandem repeat catalog and 83 human genome samples (including HG38) from English et al..^66^ Tandem repeat regions in this catalog were identified using TRF (default parameters), and the same filtering criteria were applied to establish an STR benchmark dataset based on HG38.

We further retrieved nine cancer samples from the NCBI database^70^, comprising eight lung cancer samples (DRR171429, DRR171430, DRR171431, DRR171432, DRR171433, DRR171452, DRR171453, DRR171454) and one breast cancer sample (DRR203145), and converted them into FASTQ format using the SRA Toolkit. These datasets were subsequently assembled, and assembly quality was assessed to support downstream variant detection in lung cancer. Finally, we downloaded the latest gene annotation file for T2T-CHM13v2.0 (chm13v2.0_RefSeq_Liftoff_v5.2.gff3), which corrects errors present in GENCODE v35 CAT/Liftoff and RefSeq v110 annotations and incorporates additional gene copies identified in the T2T assembly.

### Using DBSCAN and Rough Processing to Get Initial Repeats Regions

#### Anchors Density Clustering

A base pair separated by n positions is referred to as an anchor, denoted as *anchor*(*p*_1_, *p*_2_, *n*), where *p*_1_and *p*_2_represent the positions of the two flanking bases in the sequence. An STR typically consists of a dense arrangement of anchors, which may be adjacent or overlapping, forming an anchor chain that spans the entire STR and the shorter non-STR sequences at both ends. The set of all anchors separated by n positions is denoted as *A*_*n*_ = {*anchor*|*anchor*[2] = *n*}. FastSTR utilizes the DBSCAN algorithm to cluster all anchors in the set *A*_*n*_ (*n* ∈ {1,2, ⋯, 8}) from the sequence, thereby identifying all fuzzy repeat regions in the sequence.

#### Using the Distribution of Front Base Density for Initial Screening of STRs

The density of front bases effectively distinguishes non-STR sequences from true STR sequences. In FastSTR, the length of the front bases check segment is set according to the alignment score (if the alignment score is 30, the check segment length is 15; if the alignment score is 50, the check segment length is 25), and the front bases density is defined as shown in equation

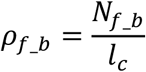

Where *l*_*c*_ refers to the length of the check segment, and *N*_*f*_*b*_ denotes the number of leading bases in a check segment. We use a sliding window to calculate the front bases density for each base, ensuring that the algorithm operates with a time complexity of *O*(*L*), where *L* is the length of the fuzzy repeats region. To determine the optimal threshold for front bases density, we randomly generated a large STR dataset and validated the effectiveness of the threshold in assessing the authenticity of STRs using a chi-square test(Supplementary Fig. 7).

#### Trivial and Obfuscation Repeat Regions Distinction

To distinguish between trivial and obfuscation repeat regions, we need to analyze the distribution of front bases’ densities in fuzzy repeat regions. FastSTR calculates the front bases’ density every 5 bp, and these values form a set *D*_*f*_*b*_. We then compute the variance *S*_*f*_*b*_ of *D*_*f*_*b*_ relative to a fixed value for a given fuzzy repeat region. If the variance is below a threshold, the region is classified as an obfuscation repeat region; otherwise, it is classified as a trivial repeat region. To determine the fixed values and thresholds, we simulated a large number of trivial and obfuscation repeat regions for various motif lengths and analyzed their distributions. Our results show that the variance of *S*_*f*_*b*_ with respect to the fixed value follows a near-normal distribution. Based on this distribution, we established thresholds such that approximately 99% of obfuscation repeat regions have an *S*_*f*_*b*_ below the threshold, and 90%-99% of trivial repeat regions have an *S*_*f*_*b*_above the threshold.

#### Context-Aware Motif Recognition Method

Based on the characteristics of STR sequences, we propose a context-aware motif recognition method. First, we traverse the STR sequence to obtain the counts and positions of all 2-mers (seeds) and 3-mers. Using the distribution of the most frequent 2-mers and the motif length, we construct a set of candidate motifs, denoted as *M*. We then apply a Tri-Gram model^85^ from the N-Gram framework to calculate the probability of each candidate motif appearing in the specified STR sequence, as described by equations.

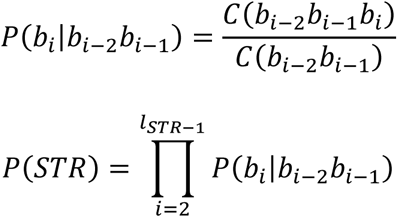

Where *C*(*b*_*i*−2_*b*_*i*−1_) denotes the frequency of the 2-mer in the given STR, and *C*(*b*_*i*−2_*b*_*i*−1_*b*_*i*_) represents the frequency of the 3-mer in the same STR. First, we calculate the probability of each 3-mer involved in the candidate motifs. Then, we multiply the probabilities of the 3-mers that comprise each motif to obtain the overall probability of the candidate motif appearing in the STR sequence. Finally, the motif with the highest probability is selected as the representative motif for the STR.

As the Tri-Gram model shows suboptimal performance for identifying STRs with motif lengths of 5-8bp, we further adopt the concept of Markov chains^86^ to analyze the feasibility of candidate motifs. Initially, we use the Tri-Gram model to filter out most unsuitable candidate motifs and construct the nucleotide transition state matrix *M*_*STR*_*base*_ by traversing the STR. Let *u*_*m*_ be the motif length and *copy*_*t*_ the theoretical copy number, so *l*_*STR*_ = *u*_*m*_ ⋅ *copy*_*t*_. Considering potential mismatches, insertions, and deletions (indels) in real STRs, we extract the two most frequent nucleotides in *M*_*STR*_*base*_ and use the Euclidean algorithm to estimate the copy number *copy*_*e*_ (as shown in equation), thereby obtaining the corrected copy number *copy*_*f*_ = *α* ⋅ *copy*_*t*_ + (1 − *α*) ⋅ *copy*_*e*_. The nucleotide transition relationship within a motif approximates a multiple of the entire STR’s transition relationship, with the multiple being *copy*_*f*_. For each retained candidate motif, we obtain its nucleotide state transition matrix *M*_*motif*_*base*_ and derive the corresponding copy number vector *copy*_*vector* using equation. It is noteworthy that when the motif length is 5 or greater, there can be instances where different candidate motifs have the same Tri-Gram probability (as shown in Construction). In such cases, it is necessary to use the Markov chain for further filtering. Similarly, when the motif length is 5 or greater, different candidate motifs may have the same *copy*_*vector* (as shown in Construction), requiring further filtering using the Tri-Gram model. Consequently, it can be proven that a combined approach using both Tri-Gram and Markov chain effectively identifies the optimal unique motif representing the STR.

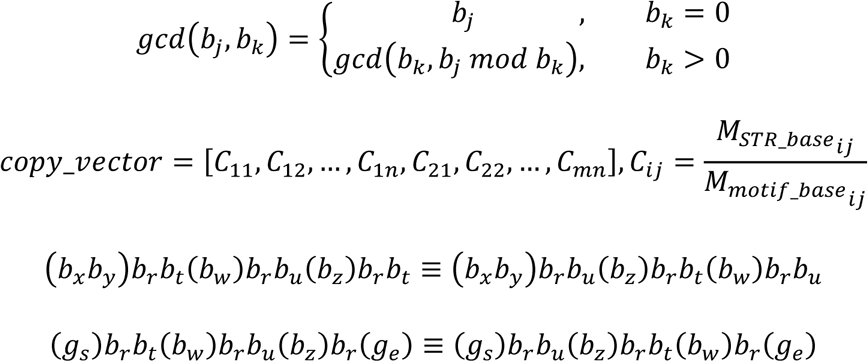

Finally, we calculate the variance of each *copy*_*vector* relative to *copy*_*f*_, considering the candidate motif with the smallest variance as the representative motif.

### Motif Mapping-Based STR Filtering Strategy

#### Signal Segment Filtering Rules for Trivial Regions

To further filter out true STRs, we map the identified motifs back to the original ambiguous repeat regions and filter out sequences that do not meet the criteria based on the mapping results. First, we construct specific regular expressions for motifs of different lengths, with longer motifs allowing more flexible matching. Next, we use the finditer function from the regex library to return segments matching the motif and assign matching scores according to predefined matching criteria. Finally, we apply a sliding window approach to identify the highest-scoring window within the ambiguous region and decide whether to retain the region based on whether the score exceeds a given threshold.

#### Obfuscation Region Partitioning Algorithm by Concentration Tests

According to the definition, true STRs and pseudo STRs within ambiguous intervals exhibit highly similar nucleotide arrangements but display significantly different motif density levels when considering exact matches. We propose the *Density Inspection Method*, which uses motif density to distinguish between true STRs and pseudo STRs. First, we design specific regular expressions for motifs of different lengths to map them back to the ambiguous repeat regions. Then, we compute the distances between adjacent mapped sites and identify potential starting points for STRs. We define the number of perfect STRs (with zero distance between mapped sites) within an inspection segment of approximately 75 bp as *C*_*perfect*_ ≈ 75 ∕ *u*_*m*_. Using a sliding window approach, we calculate the average distance between consecutive *C*_*perfect*_mapped points, and determine if a segment is a true STR based on whether the average is below a set threshold. The threshold is established by evaluating the performance of the Density Inspection Method on a large, randomly generated dataset of true and pseudo STRs.

#### Probe Alignment-Based Approximate Boundary Correction

After initial filtering and segmentation, the ends of the preliminary STRs may still contain non-STR sequences, necessitating further trimming and refinement. Based on the motif and recognition criteria, an STR probe of length *l*_*p*_ is constructed (e.g., *l*_*p*_ is 15 for an alignment score of 30, and 25 for a score of 50). A local sequence alignment algorithm is used to slide the probe inward from both ends of the initial STR until the alignment score reaches the threshold, at which point the sliding stops. To enhance filtering efficiency, the probe advances in 5bp increments. If the sliding does not stop and the entire initial STR is traversed, that STR is discarded.

### Candidate STRs Merging Rule

The final step before sequence alignment is to merge candidate STRs. To define the merging criteria, we first performed density clustering on the anchor points of all STRs identified by TRF in the human genome, obtaining all STRs disrupted due to clustering, denoted as *I*_*STR*_. Analysis revealed that the proportion of STRs in *I*_*STR*_ relative to all STRs on each chromosome is generally within 0.5% (see Supplementary Figure 6), and over 99% of the STRs in *I*_*STR*_ have disruption lengths within 100bp. Based on this, we established the merging criteria: two candidate STRs are considered for merging if their motifs are similar and their distance is within 100bp. Given that the disrupted sequences typically lack significant STR structures, we further assess the relationship between the candidate STRs and the consensus motif for those meeting the merging criteria. To simplify the merging process, we prioritize accepting longer STRs. If the motifs of the candidate STRs match the consensus motif, they are directly merged, with additional capture operations implemented to correct potential mismerging. For candidate STRs with differing motifs, signal filtering and probe alignment are applied to determine whether the consensus motif aligns with their structure, guiding the decision to merge.

### High-Speed Segmented Global Sequence Alignment for Ultra-Long STRs

The merged candidate STRs were analyzed using the local sequence alignment algorithm (Smith-Waterman algorithm) to calculate their match percentage (*P*_*Match*_), indel percentage (*P*_*Indel*_), and alignment score (*S*_*align*_). Let *N*_*mismatc*ℎ_, *N*_*insert*_, and *N*_*delete*_ represent the number of mismatched bases, inserted bases, and deleted bases in the aligned STRs, respectively. The definitions of *P*_*Match*_ and *P*_*Indel*_ are shown in formulas and. Assuming the length of the reference STR corresponding to the STR being aligned is 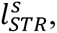 the traditional local sequence alignment algorithm involves iterating over all base pairs (*i*, *j*), with its time complexity is 0(*l*_*STR*_^2^). For STRs with complex structures, the time complexity increases exponentially.

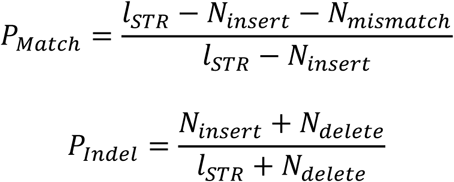

To address this, we developed a high-speed alignment algorithm tailored for ultra-long STRs—Segmented Global Sequence Alignment Algorithm. This algorithm achieves rapid sequence alignment while maintaining results comparable to those of local sequence alignment. It comprises two core modules: Real-Time Path Tracking Alignment Algorithm and Motif-Constrained Global Alignment Algorithm. Initially, the motif length of the STR is used to define the marker motif length, and the finditer function is employed to identify all marker motif positions. Overlapping positions within chaotic sequences—where the optimal matching position cannot be determined—are treated as invalid matches. This step allows the majority of the ultra-long STR to be efficiently aligned, leaving only a small number of unaligned fragments for further analysis. If unaligned fragments are present at the starting end, the Motif-Constrained Local Alignment Algorithm is used to determine the optimal starting position while ensuring the distance between this position and the fragment endpoint is a multiple of the motif length. For other unaligned fragments (including chaotic sequences), the Real-Time Path Tracking Alignment Algorithm is applied to find the optimal alignment. Notably, the real-time path tracking algorithm performs global sequence alignment, ensuring the entire sequence of intermediate fragments is included in the alignment. The best alignment result is then determined based on alignment scores at motif-multiple positions. To efficiently trace back alignment paths, the real-time path tracking algorithm leverages the iterative nature of optimal matches/mismatches in sequence alignment, introducing an additional dynamic programming step to construct the count of matches and mismatches along the optimal alignment path. Based on the two rounds of dynamic programming, we propose a Normalized Geometric Path Quality Metric for Alignment, which can calculate match percentage, indel percentage, and alignment scores in linear time complexity.

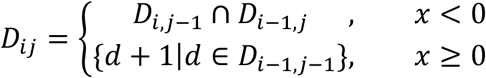

To determine the marker motif lengths corresponding to different motif lengths and the segmented global sequence alignment parameters corresponding to various local sequence alignment parameters, we randomly generated 3,000 ultra-long STRs with lengths ranging from 7,000 to 70,000. These sequences were analyzed using both the Smith-Waterman algorithm and the Segmented Global Sequence Alignment Algorithm(Supplementary Fig. 9).

### STRs Merging and Overlap Handling Strategy

#### Large-Scale Merging Strategy for STRs in Adjacent Sub-Reads

After identifying all STRs on a single sub-read, the next step is to merge overlapping STRs from adjacent sub-reads to construct consensus STRs. The overlapping relationships between STRs fall into exactly four types: left overlap, right overlap, full overlap, and mutual embedding. It is guaranteed that each overlapping relationship involves exactly two STRs, one on the preceding sub-read and the other on the following sub-read, which significantly reduces the complexity of constructing consensus STRs. Additionally, we analyzed the distribution of ultra-long STRs across multiple sub-reads and proposed rules for merging overlapping STR segments into a complete ultra-long STR. Specifically, when STRs on multiple adjacent sub-reads share the same motif and fully cover their respective sub-reads, we create an array to store these overlapping STRs, which are then merged into a single complete consensus STR.

#### Overlap STRs Selection Strategy Based on Interval Gain Decision Algorithm

The final step of FastSTR addresses the selection of overlapping STRs. Since there is currently no unified standard for handling different STRs identified in the same region, we aim to retain all STRs that may have biological significance, removing only those with short lengths and low quality due to erroneous motif identification. To this end, we propose an overlap STR selection strategy based on an interval gain decision algorithm, which emulates decision-making behavior by analyzing whether a specific STR in a given region is worth exploring to determine if it should be retained.

Specifically, we first define the conditions for overlapping relationships and mark the regions covered by STRs that satisfy these conditions. Next, we define the mining cost 𝜛_*c*_ as the number of edits required to convert a marked region into a specific STR, the mining gain 𝜛_*p*_ as the standard STR length 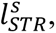 and the net mining gain 𝜛_*g*_ = 𝜛_*p*_ − 𝜛_*c*_. Let the length of the marked region be *L*_*mark*_; 𝜛_*c*_ is calculated using formula (12). To simplify the calculation, we assume the insertion and deletion base counts are approximately equal. Combining formulas (8) and (9), we derive formula (13).

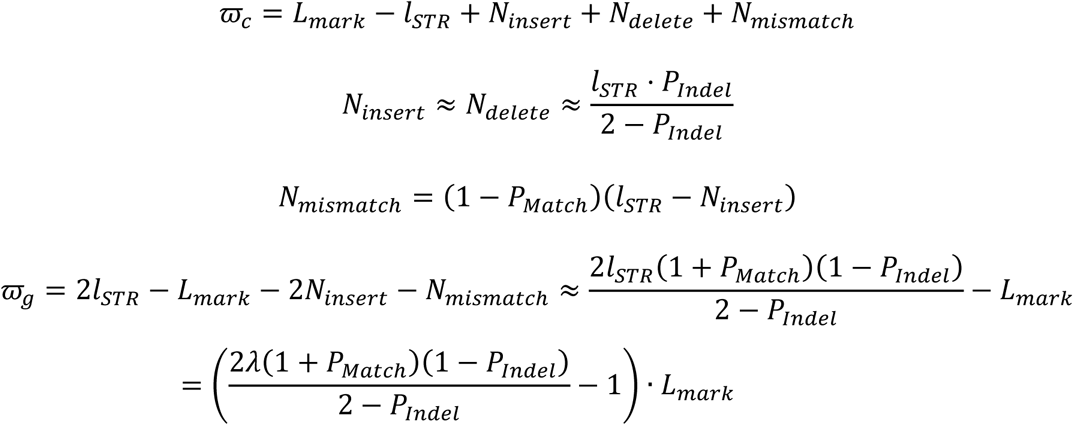

Furthermore, since 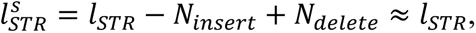 we define the proportion of the STR length to the marked region length as *λ*, leading to the final expression for 𝜛_*g*_ in formula (14). Finally, we determine whether to retain the STR based on whether 𝜛_*g*_ is non-negative. For example, when *P*_*Match*_ = 0.8 and *P*_*Indel*_ = 0.15, an STR must account for at least 60% of the marked region length to be retained. Notably, whether 𝜛_*g*_ exceeds zero depends solely on the STR’s *P*_*Match*_, *P*_*Indel*_, and length, avoiding the pitfalls of subjective judgment in selecting overlapping STRs.

### Comprehensive STR catalogue construction and polymorphism analysis using FastSTR

We applied FastSTR (default settings) to the T2T reference genome to identify STRs. To mitigate the impact of overlapping STR intervals, we merged overlapping regions into contiguous intervals using bedtools^65^ merge. Overlaps among the T2T annotation, TRF results, and FastSTR calls were then quantified with bedtools^65^ intersect, adopting the criterion that any ≥1 bp overlap constitutes a valid intersection. As overlap counts can vary depending on input order, we evaluated all permutations and retained the minimum value as the final count.

We next characterized the motif composition of STRs uniquely detected by FastSTR and those overlapping with the T2T annotation. Using custom scripts, we further quantified STR counts per 1-Mb interval along each chromosome, based on sequence counts rather than interval counts. To validate newly identified STRs in centromeric regions, we extracted satellite sequences from the T2T tandem repeat annotation, filtered for HSATII, and profiled its distribution across centromeres of chromosomes 1 and 16. Newly discovered STRs were exported in BED format and visualized in the UCSC Genome Browser, including HSATII in chromosomes 1 and 16 and HSAT1B in the Y chromosome.

To generate a comprehensive hg38 STR benchmark, we obtained the adotto_TRregions_v1.2.1.bed catalog from English et al.^66^, which aggregates tandem-repeat annotations from multiple sources. Since the original analysis applied a TRF score threshold of 5 (much lower than the default of 50), many low-quality repeats shorter than 25 bp were included. We therefore re-annotated all TR regions using TRF (parameters “2 5 7”) and filtered for STRs with motif lengths ≤8 bp to build an improved hg38 STR benchmark.

We analyzed 82 haploid assemblies with FastSTR (default settings), generating STR BED files via custom scripts. Each assembly was aligned to the T2T and hg38 references with minimap2, and STR coordinates were lifted over to both references using paftools liftover. Variants were called with paftools call, and we defined STR variants as indels ≥1 bp occurring within STR loci, with a minimum quality score of 20. Resulting variants were classified into four categories: (i) variants present in both the T2T annotation and hg38 benchmark, (ii) variants unique to T2T, (iii) variants unique to hg38, and (iv) novel STR variants.

To assess similarity between assemblies, we calculated the Jaccard index as the ratio of intersecting STR variant intervals to their union. To avoid bias from overlapping variants, intervals were first merged using bedtools merge, with the option “-c 4 -o collapse” applied to retain interval labels and track which assemblies overlapped each locus. Using these labels and population metadata, we identified population-specific STR variant loci (shared by multiple assemblies within the same population) and individual-specific STR variant loci (unique to a single assembly).

### Detection of Novel Pathogenic STR Variants in Lung Cancer

To systematically investigate STR variation, we generated high-quality genome assemblies for nine cancer samples (eight lung cancer and one breast cancer). Raw sequencing reads were quality-controlled with Chopper^87^, retaining those with quality scores ≥ 7 and lengths ≥ 5 kb. Assemblies were constructed using Flye^88^ and polished through multiple rounds of Medaka^89^ correction. Genome completeness and assembly quality were assessed with BUSCO^90^ and QUAST^91^.

After assembly and polishing, STR loci in each tumor genome were identified using FastSTR with default parameters, and per-sample STR BED files were generated with custom scripts. The nine tumor assemblies were aligned to the T2T reference genome with Minimap2, and STR loci were mapped with paftools. Following the same procedure, STR variants were extracted for each sample. To minimize biases caused by overlapping intervals, we merged STR variant regions from lung cancer and normal samples using bedtools merge, and quantified overlaps among normal, lung cancer, and breast cancer samples. Based on STR and STR variant BED files, as well as assembly data, we computed the distribution of STRs and STR variants per megabase across tumor and normal samples using custom scripts.

To determine functional localization, BED files for genomic features were generated from the official T2T annotation (chm13v2.0_RefSeq_Liftoff_v5.2.gff3), prioritized in the order: CDS, UTR, intron, ncRNA, 1 kb upstream of transcription start sites (TSS_up1k), and intergenic regions. STR variants from normal, lung cancer, and breast cancer samples were intersected with these regions using bedtools, and enrichment of STR variants in lung cancer samples was evaluated with Fisher’s exact test.

We next extracted STR variants occurring exclusively in lung cancer samples and classified them by genomic context (CDS, UTR, intron, ncRNA, TSS_up1k, intergenic). Genes associated with STR variants located in CDS, UTR, or TSS_up1k across eight lung cancer samples were subjected to GO and KEGG enrichment analysis using g:Profiler (https://biit.cs.ut.ee/gprofiler/gost) ^92^. Benjamini–Hochberg FDR correction was applied, with all other parameters set to default. Only terms with sizes between 10 and 500 were retained, consistent with the default settings of the clusterProfiler package^93^.

Finally, we selected genes that exhibited both high coverage across the eight lung cancer samples and significant enrichment in GO analysis, and visualized their STR variation using the UCSC Genome Browser. Corresponding reference (T2T) and sample sequences at these loci were retrieved, and amino acid–level variation was analyzed using ANNOVAR^94^.

### Evaluation

To comprehensively benchmark FastSTR against four widely used STR detection tools—Tandem Repeats Finder (TRF), mreps, TRASH, and T-reks—we ran each program on the same reference genomes using default or developer-recommended parameters to ensure a fair comparison. Because TRF is broadly recognized as the gold-standard “ground truth,” its output was used as the reference for assessing other tools’ performance in sensitivity (recall), precision, F₁ score, interval boundary error, and motif identification specificity. Runtime and peak memory usage were recorded for all tools.

We quantified STR counts based on the number of distinct motifs reported by TRF, rather than the number of intervals, to capture the biological significance of different motifs within the same genomic region. Given that TRF reports only the top three scoring repeats per interval—potentially omitting valid STRs—any STR not falling within a TRF-annotated region was classified as a false positive. To align with TRF’s minimum detectable length, we filtered each tool’s output to include only STRs ≥ 25 bp. We considered an STR detection valid if it overlapped the ground truth by at least 1 bp, and defined:

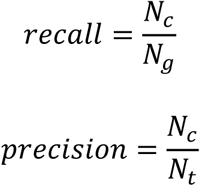

where *N*_*g*_ is the total number of STRs identified by TRF, *N*_*t*_ is the count of STRs reported by the evaluated tool, and *N*_*c*_ is the number of tool-reported STRs that validly cover the ground truth. We also analyzed the distribution of boundary deviations for each validly detected STR.

To evaluate motif-level accuracy, we randomly selected 5000 ground-truth STRs on human chromosome 1 and compared the best-match motifs predicted by each tool against the true sequences. To evaluate the accuracy of FastSTR in identifying STR copy numbers, we retrieved all STRs detected by FastSTR across each species. To ensure a fair comparison of copy number estimates, we filtered the results to retain only STRs whose motifs matched those identified by TRF and whose genomic overlap was ≥15 bp. We then compared the copy number differences between the two methods. Since mreps and T-reks lack native parallelization, we split each genome by chromosome and processed them in parallel, summing the per-chromosome runtimes and taking the maximum per-chromosome memory usage. FastSTR and TRASH were both executed on a 72-core Intel® Xeon® Platinum 8275CL (3.00 GHz) cluster, with memory profiling performed via the smem utility.

## Reporting summary

Further information on research design is available in the Nature Portfolio Reporting Summary linked to this article.

## Data availability

## Code availability

https://github.com/XL-BioGroup/FastSTR

## Acknowledgements

This work was supported by the National Natural Science Foundation of China (No.62433016,No.62002388); The Fundamental Research Funds for the Central Universities (No.D5000230363); King Abdullah University of Science and Technology (KAUST) Office of Sponsored Research (OSR) under award numbers FCC/1/1976-44-01, FCC/1/1976-45-01, REI/1/5202-01-01, REI/1/4940-01-01, and RGC/3/4816-01-01.

## Author contributions

X.L. and L.W. conceived of the presented idea. L.W., X.L. and B.Z. designed the computational framework and analyzed the data. X.L., L.W., M.J., X.L.,B.C., B.Z., and X.S. conducted the clinical evaluation. X.L. provided valuable intellectual input during the software refinement process. X.G., B.Z. and X.S., supervised the findings of this work. X.L., L.W., B.Z., X.S., and X.G. took the lead in writing the manuscript and supplementary information. All authors discussed the results and contributed to the final manuscript.

## Competing interests

The authors declare no competing interests.

## Additional information

**Supplementary information**

**Correspondence and requests for materials**

**Peer review information**

**Reprints and permissions information**

## Notes

### Competing Interest Statement

The authors have declared no competing interest.

