## Supplementary Information for "Efficient Identification of Short Tandem Repeats via Context-Aware Motif Discovery and Ultra-Fast Sequence Alignment"

#### Supplementary Methods

##### Principle of Copy Number Estimation Based on the Euclidean Algorithm

In this method, we start from the assumption that a short tandem repeat (STR) is composed of multiple highly similar motif repeats. Consequently, the relative frequency of each base within a motif is proportionally amplified across the entire STR. Leveraging this property, the approximate copy number relative to the motif length can be inferred from the observed base counts of the STR.

Let the motif length be $u_m$ and the total length of the STR be $l_{\mathrm{STR}}$. Ideally, $l_{\mathrm{STR}} = u_m \cdot \mathrm{copy}_t$, where $\mathrm{copy}_t$ denotes the theoretical copy number. Define the multiplicity vector of bases within a motif as $\boldsymbol{\rho} = (q_1, q_2, \ldots, q_r)$, where $q_i$ represents the number of occurrences of the $i$-th base in one motif, satisfying $\sum_i q_i = u_m$. In the observed STR, the count of the $i$-th base is denoted as $b_i$. Under substitution, insertion, or deletion noise, we approximate:

$$b_i = q_i \cdot \mathrm{copy}_t + \varepsilon_i,$$

where $\varepsilon_i$ represents small perturbations introduced by sequencing or alignment errors. Since $b_i$ is approximately proportional to the product of $q_i$ and $\mathrm{copy}_t$, the greatest common divisor (GCD) property of integers can be exploited for estimation. For several counts not dominated by noise, the following approximate relation holds:

$$\gcd(b_i, b_j, \ldots) \approx \mathrm{copy}_t \cdot \gcd(q_i, q_j, \ldots)$$

Therefore, if $\gcd(q_i, q_j, \ldots)$ can be estimated or is known, $\mathrm{copy}_t$ can be approximately recovered.

In practical implementation, we select the two or several most frequent bases (excluding those with very small counts likely caused by noise) and compute their GCD using the Euclidean algorithm to obtain an estimated copy number $\mathrm{copy}_e$.

Before computing the GCD, a simple noise filtering step is applied: base counts much smaller than the second largest count (likely caused by single substitutions or insertions) are discarded. The specific handling strategies are as follows:

1) Motif contains two base types, one with multiplicity of 1

Let $\boldsymbol{\rho} = (1, q_{\text{maj}})$. In this case, the second most abundant base (denoted $b_{\text{maj}}$) approximately equals $1 \cdot \text{copy}_t$. Because substitution/insertion noise is relatively minor, we directly take the second largest count as an approximate value of the copy number, i.e., $\text{copy}_e = b_{\text{maj}}$.

2) Motif contains two base types, both with multiplicities > 1

Let $\boldsymbol{\rho} = (a, b)$ with $\min(a, b) \geq 2$. Both base counts scale with their respective multiplicities and the theoretical copy number. If $a$and $b$are unequal but share a common divisor $g = \text{gcd}(a, b) > 1$, then:

$$\text{gcd}(b_1, b_2) \approx g \cdot \text{copy}_t.$$

Thus, we compute $g_b = \text{gcd}(b_1, b_2)$ and divide it by $g$ (or by the known multiplicity $a$or $b$) to obtain $\text{copy}_t \approx g_b/g$. If $a = b$ (e.g., (3,3)), then $\text{copy}_t \approx b_1/a$(or $b_2/a$).

3) Motif contains more than two base types

In most practical cases, the two most frequent bases suffice for analysis. If the internal multiplicities of these two bases are coprime ($\text{gcd}(q_{(1)}, q_{(2)}) = 1$), the estimated copy number is directly given by:

$$\text{copy}_e \approx \text{gcd}(b_{(1)}, b_{(2)}).$$

If their multiplicities are equal or have a multiple relationship ($\text{gcd}(q_{(1)}, q_{(2)}) > 1$), normalization is applied following the rule described in case (2). When more bases are available, the estimation can be extended by computing the GCD among the most frequent bases and normalizing by their common divisor within the motif.

Finally, a weighted fusion is used to reduce single-source bias:

$$\text{copy}_f = \alpha \cdot \text{copy}_t + (1-\alpha)\cdot \text{copy}_e,$$

where $\alpha \in [0,1]$ is a weighting coefficient that can be adjusted according to sequence quality, noise level, or prior confidence in $\text{copy}_t$ and $\text{copy}_e$.

#### Principle of Unique Motif Identification Using Tri-Gram and Markov Chain Representation

Let $s = s_1 s_2 \cdots s_L$ denote a DNA motif (or nucleotide sequence), where $L \geq 5$. We introduce the following definitions.

**Definition 1**: Tri-Gram Set

The Tri-Gram set $T(s)$ is defined as the unordered set of all distinct substrings of length three obtained from a sliding window across $s$:

$$T(s) = \{\, s_i s_{i+1} s_{i+2} \mid 1 \leq i \leq L-2 \,\}.$$

For example, if $s = \text{ATGCA}$, then

$$T(s) = \{\text{ATG}, \text{TGC}, \text{GCA}\}.$$

**Definition 2**: Markov Chain Representation

The Markov chain representation of the motif corresponds to the ordered transition sequence:

$$M(s) = s_1 \to s_2 \to \cdots \to s_L,$$

which preserves the full sequential order (i.e., the transition structure between adjacent nucleotides).

In the following, we first discuss the information contained only in the Tri-Gram set $T(s)$. By applying graph-theoretic reasoning, $T(s)$ can be mapped to a directed graph, and we analyze whether this graph uniquely determines the original linear sequence $s$.

A unique directed graph $G = (V, E)$ can be constructed from $T(s)$ as follows:

**Definition 3:** Vertex set $V$: All ordered 2-mers (length-2 substrings) appearing in $s$:

$$V = \{ s_i s_{i+1} \mid 1 \leq i \leq L - 1 \}.$$

Note that order matters; for example, "AG" and "GA" are treated as distinct vertices.

**Definition 4:** Edge set $E$: For each tri-nucleotide $t = xyz \in T(s)$ (where $x, y, z$ are individual bases), add a directed edge:

$$e: (xy) \rightarrow (yz).$$

Each directed edge therefore corresponds one-to-one to a 3-mer labeled by $xyz$.

This construction is essentially a De Bruijn-type representation with $k = 3$, expressed using vertices of order 2. Hence, the Tri-Gram set $T(s)$ uniquely determines the graph $G$, since each 3-mer uniquely defines a directed edge from its prefix 2-mer to its suffix 2-mer.

**Lemma 1** (No Parallel Edges)

In the directed graph $G$ constructed from $T(s)$, no two distinct parallel directed edges exist between the same ordered pair of vertices.

**Proof**.

Assume there exist two distinct edges $u \rightarrow v$ and $u \rightarrow v$. Each edge corresponds to a distinct 3-mer. However, each edge is uniquely determined by its endpoints: if $u = xy$ and $v = yz$, then the corresponding 3-mer must be $xyz$. Thus, two distinct edges connecting the same pair of vertices cannot exist.

**Lemma 2** (Minimum Length Constraint for Branch–Merge Structures)

If the graph $G$contains a branch–merge structure—that is, there exist two distinct directed paths from vertex $u$to vertex $v$with no overlapping internal vertices—and we expect such a structure to yield *distinct feasible reconstructions* of motifs of length at least 5 (i.e., distinct linear sequences $s$and $s'$sharing the same Tri-Gram set), then the segment between $u$and $v$(measured in edge count) must contain at least three edges.

In other words, if the two different $u$-to-$v$ paths each contain ≤ 2 edges, no distinct reconstructions of length ≥ 5 can exist.

**Proof** (by contradiction, case analysis).

Let there exist two distinct paths $P$ and $Q$ from $u$ to $v$.

1) If the path length equals 1 (i.e., $u \to v$ appears twice), then there would be two parallel edges between $u$ and $v$, contradicting **Lemma 1**. Hence, the length cannot be 1.
2) If the path length equals 2 (i.e., $u \to p \to v$ and $u \to q \to v$ with $p \neq q$), denote $u = xy$. By the edge construction rule, the first base of $p$ must be $y$, thus $p = ya$ (for some base $a$); similarly, $q = yb$. Since edges $p \to v$ and $q \to v$ exist, the first base of $v$ must be $a$ and $b$ respectively, i.e., $v = ac$ and $v = bc$. From $ac = bc$, we obtain $a = b$, hence $p = q$, contradicting $p \neq q$.

Therefore, for distinct linear reconstructions (of length ≥ 5) to exist, the branch–merge segment must contain at least three edges.

**Remark**.

Intuitively, a motif of length $k$ corresponds to a path of $k - 2$ edges in the graph. When only the unordered Tri-Gram set is retained (i.e., only the existence of edges is preserved, while edge multiplicities and starting positions are discarded), two distinct edge sequences (representing distinct linear motifs) can coexist without altering the edge set *only* if the branch–merge region is sufficiently long (≥ 3 edges). This explains why ambiguity in sequence reconstruction arises only beyond this threshold.

**Lemma 3** (Self-Loops Correspond to Triple Identical Bases)

If the vertex $XX$ in graph $G$ has a self-loop—i.e., there exists an edge $(XX) \to (XX)$—then the corresponding 3-mer must be $XXX$.

**Proof**.

A self-loop implies the existence of a 3-mer $xyz$ such that the prefix 2-mer $xy = XX$ and the suffix 2-mer $yz = XX$. This requires $x = X$, $y = X$, and $z = X$, hence the 3-mer is $XXX$.

Finally, for a motif $s = s_1 s_2 \cdots s_k$, the corresponding directed graph $G = (V, E)$ satisfies:

$$|V| = k - 1, |E| = k - 2.$$

In particular, when $k \in [5,8]$, we have $|E| \in [3,6]$.

In the directed graph $G$ uniquely determined by the Tri-Gram set $T(s)$, every directed walk from the start vertex to the end vertex has a length equal to $|E|$. If two distinct motifs $s$ and $s'$ share the same Tri-Gram set, i.e. $T(s) = T(s')$, then they correspond to the same directed graph $G$. In that case, $G$ must contain at least two distinct vertex-connected directed walks $W_1, W_2$, each of which traverses every edge in $E(G)$.

In other words, $T(s)$ is non-unique $\Rightarrow$ $G$ admits multiple Euler-type walks, where an *Euler-type walk* refers to a directed walk that visits every edge exactly once. If a directed graph $G$ possesses at least two distinct vertex-connected walks that both cover all edges $E(G)$, then the topology of $G$ must belong to one of the following categories:

1) Simple cycle: all vertices have in-degree and out-degree equal to 1.
2) Branching path: $G$ contains a branching subpath that forms part of a simple cycle.
3) Self-loop: $G$ contains a self-loop that belongs to a simple cycle.
4) Parallel edges: $G$ contains multiple directed edges connecting the same pair of vertices.
5) Combinations of the above structures.

From **Lemma 1** (absence of parallel edges), case (4) cannot occur. We therefore discuss cases (1)–(3) in order.

Case (1): Simple Cycle Structure

**Theorem 1** (Simple cycles yield multiple Euler-type walks)

If $G$ is a simple directed cycle—i.e., every vertex has exactly one incoming and one outgoing edge—and $|E(G)| \geq 3$, then $G$ admits multiple vertex-connected Euler-type walks. Consequently, the corresponding motif is not unique.

**Proof**.

Let the graph $G$ consist of a vertex set $V = \{v_1, v_2, \ldots, v_n\}$ and an edge set $E = \{(v_i, v_{i+1}) \mid 1 \leq i \leq n-1\} \cup \{(v_n, v_1)\}$. Because each vertex has both in-degree = 1 and out-degree = 1, any vertex on the cycle can serve as a starting point to generate a distinct cyclic sequence:

$$v_i \rightarrow v_{i+1} \rightarrow \cdots \rightarrow v_n \rightarrow v_1 \rightarrow \cdots \rightarrow v_{i-1}.$$

These correspond to *cyclic shifts* of the motif, denoted as $\mathrm{Rot}_i(s)$. Therefore, a simple cycle necessarily gives rise to multiple equivalent Euler-type walks, each representing a rotation of the same motif. When $\mid E \mid = 2$, the graph degenerates into a single self-loop, corresponding to a motif of length < 5, which is outside the scope of consideration. For example, the motif AGCAG produces a 3-vertex simple cycle graph $G$ with both in-degree and out-degree = 1 **(Supplementary Fig.1 a)**. Different traversal starting points correspond to linear sequences such as AGCAG, GCAGA, and CAGAG, which share the same Tri-Gram set.

Case (2): Trivial Cycle with a Branch

**Theorem 2** (Motif Length Constraint for Branched Cyclic Structures)

If a directed graph $G$ contains a branching path (i.e., a vertex with out-degree ≥ 2 or in-degree ≥ 2), and this branching path is a subgraph of a trivial cycle, then the minimum motif length required to produce distinct motifs is 8 bp.

**Proof**.

According to **Lemma 2**, in order to form two *distinct* Euler-type walks, the branch–merge segment must contain at least three edges. A complete trivial cycle itself contains at least three edges, so after including the branching segment, the total number of edges must satisfy:

$$\mid E(G) \mid \geq 3 + 3 = 6$$

By Proposition, we have $\mid E(G) \mid = k - 2$, thus $k \geq 8$. When $k = 8$, there exists a minimal example satisfying this condition. For instance:

$$\mathrm{motif}_1 = \mathrm{AGCAGTAG}, \mathrm{motif}_2 = \mathrm{AGTAGCAG},$$

These two motifs share the same Tri-Gram set, yet exhibit different linear orders **(Supplementary Fig.1 b)**.

Case (3): Trivial Cycle Containing Self-Loops

**Theorem 3** (Combination of Self-Loops and Trivial Cycles)

If a directed graph $G$ contains one or more self-loops, and these self-loops are part of a trivial cycle, then:

1) If there is only one self-loop, all possible walks generate motifs that are equivalent under cyclic rotation.
2) If there are two or more self-loops, non-equivalent motifs can occur only when the motif length is 8 bp.

**Proof**.

1) Single self-loop.

Let the trivial cycle have length $n \geq 3$. If there exists a single vertex $v_i = XX$ with a self-loop, then any Euler-type walk starting from that vertex or any other vertex on the cycle generates motifs that differ only by a cyclic shift, and no new non-equivalent linear sequence can be formed. For example:

$$\text{AACAAA} \Rightarrow \text{AAAAAC},$$

which are equivalent under cyclic rotation.

2) Two self-loops.

When the cycle length is 3 or 4, excessive repetition of bases causes all walks to remain equivalent. However, when the cycle length is 4 and two self-loops are present, the total number of edges is $|E| = 6$, corresponding to a motif length of $k = 8$. In this case, the different arrangements of the two self-loops yield two distinct walks, for example:

$$\text{AAACCCAA,CCCAAACC},$$

which share the same Tri-Gram set **(Supplementary Fig.1 c)**, yet differ in base order, thus constituting a genuine case of non-unique reconstruction.

From the previous analysis, we have established that if two distinct motifs $s$ and

$s'$ share the same Tri-Gram set, $T(s) = T(s')$, then they correspond to the same directed graph $G$, and $G$ must contain at least two distinct vertex-connected walks. These non-unique scenarios include:

1) Simple cycles;
2) Simple cycles with branching paths (corresponding to motif length $k = 8$);
3) Simple cycles containing self-loops (also corresponding to $k = 8$).

We now rigorously prove that in all these cases, the Markov chain representations $M(s)$ and $M(s')$ are necessarily distinct. Therefore, by combining the Tri-Gram set $T(s)$ and the Markov chain $M(s)$, a motif can be uniquely identified.

For any motif

$$s = s_0 s_1 s_2 \cdots s_n,$$

its corresponding Markov chain is

$$M(s) = s_0 \to s_1 \to s_2 \to \cdots \to s_n,$$

which can be viewed as a directed graph $G_M$ whose vertices correspond to the nucleotide alphabet $\Sigma = \{A, T, G, C\}$ and whose edges represent transitions between adjacent bases. Clearly, if two motifs differ in the first or last base, their corresponding Markov chain graphs have different start or end states.

**Lemma 4** (Uniqueness of Start and End States)

Let the Markov chain of a string $s = s_0 s_1 \cdots s_n$ be

$$M(s) = s_0 \to s_1 \to \cdots \to s_n.$$

If another string $s' \neq s$ satisfies $s'_0 \neq s_0$ or $s'_n \neq s_n$, then

$$M(s') \neq M(s).$$

**Proof**.

Construct a directed graph $G_M$ with all characters as vertices. Except for the start $s_0$ and end $s_n$, each intermediate vertex $s_i (1 \leq i \leq n-1)$ has equal in-degree and out-degree. By Eulerian path theory, a graph satisfying

$$\sum_{v \in V} | \deg^+(v) - \deg^-(v) | = 2$$

has an Eulerian path uniquely determined by its start and end vertices. Hence, any difference in the first or last base guarantees $M(s) \neq M(s')$.

**Theorem 4** (Markov Chain Distinctness for Simple Cycles)

If two distinct motifs $s$ and $s'$correspond to the same simple cycle graph $G$, then their Markov chains are necessarily distinct.

**Proof**.

All vertices (2-mers) in a simple cycle are distinct.Different motifs correspond to different starting points on the same cycle. Since their first and last bases differ, **Lemma 4** guarantees $M(s) \neq M(s')$.

**Theorem 5** (Markov Chain Uniqueness for Branched Simple Cycles)

If two 8-bp motifs $s$ and $s'$share the same Tri-Gram set $T(s) = T(s')$, and their directed graph $G$ is a simple cycle containing a branching path, then under a fixed lexicographic order, their Markov chains are distinct.

**Proof**.

Let the two distinct branch paths traverse vertex sequences:

$$P_1: X\,Y\,P\,X\,Y\,Q\,X\,Y, P_2: X\,Y\,Q\,X\,Y\,P\,X\,Y,$$

where $X, Y, P, Q$ are nucleotides with $P \neq Q$.

Their Tri-Gram sets are identical because each 3-mer appears in both paths. However, expanding in lexicographic order (e.g., using $XY$ as the initial 2-mer) yields:

$$s = XYXYPXYQ, s' = XYXYQXYP,$$

with Markov chains:

$$M(s): X \to Y \to X \to Y \to P \to X \to Y \to Q,$$

$$M(s'): X \to Y \to X \to Y \to Q \to X \to Y \to P.$$

Since $P \neq Q$, it follows that $M(s) \neq M(s')$.

**Theorem 6** (Markov Chain Uniqueness for Simple Cycles Containing Self-Loops)

If two 8-bp motifs $s$and $s'$correspond to the same graph $G$ containing self-loops, then their Markov chains are distinct.

**Proof**.

In a simple cycle with self-loops, different walks may correspond to distinct motifs, but the traversal order of self-loops ensures differing first or last bases. For example:

$$s_1 = AAACCCAA, s_2 = CCCAAACC.$$

Although their Tri-Gram sets are identical, their Markov chains are:

$$M(s_1): A \to A \to A \to C \to C \to C \to A \to A,$$

$$M(s_2): C \to C \to C \to A \to A \to A \to C \to C.$$

The differing start and end states guarantee $M(s_1) \neq M(s_2)$.

**Theorem 7** (Uniqueness Theorem)

For any motif of length $k \geq 5$, if there exists $s' \neq s$such that $T(s') = T(s)$, then necessarily

$$M(s') \neq M(s).$$

In other words, no two distinct motifs share both the same Tri-Gram set and the same Markov chain representation.

**Proof**.

From Theorems 7–9, all graph structures that could produce $T(s) = T(s')$ (simple cycles, branched cycles, or cycles with self-loops) result in differing start/end bases or transition sequences. By **Lemma 4**, differing start or end bases ensure distinct Markov chains. Therefore, it is impossible for $T(s) = T(s')$ and $M(s) = M(s')$ to hold simultaneously.

By **Theorem 7**, for any short tandem repeat (STR) motif of length $k \geq 5$, the pair

$$(T(s), M(s))$$

constitutes a fully discriminative feature set. For any two distinct motifs $s \neq s'$, at least one of the following holds:

- $T(s) \neq T(s')$, or
- $M(s) \neq M(s')$.

Hence, the combined information of Tri-Gram sets and Markov chains uniquely determines the linear structure of a motif.

**Principle of Efficient Alignment in the Segmented Global Sequence Alignment Algorithm**

We first introduce the necessary definitions:

1) Let the motif (repeating unit) be denoted as $t$, with length $l$ (bp).
2) The reference sequence (perfect repeat) is

$$R = t^c = (\underbrace{t\,t\,\cdots\,t}_{c\text{ copies}}),$$

with total length $\mid R \mid = n = c \cdot l$.

3) The observed sequence (or read) is denoted by $Q$, with length $\mid Q \mid = m$.
4) Alignment uses a global sequence alignment scoring model with simple linear gap penalties, with scoring parameters:
   a) match score $\alpha$,
   b) mismatch score $\beta$,
   c) gap penalty $\gamma$ per gap (insertion or deletion), usually $\alpha > 0$ and $\beta, \gamma < 0$.

The core task is to prove that under certain biological/model assumptions, aligning only the blocks that differ from the motif $t$, while "skipping/reusing" identical blocks, yields a segmented global alignment whose statistics (number of matches, mismatches, insertions, deletions) are equivalent or highly consistent with those obtained by a standard full-length global alignment. Furthermore, these counts can be obtained directly in a single dynamic programming pass without backtracking.

**Assumptions** (Local Errors and Gap Constraints)

**A1.** Locality**.** Variations in the observed sequence $Q$ relative to the reference $R = t^c$ occur only within individual motif copies; i.e., each substitution or indel falls entirely within a single copy of $t$. No large-scale structural misalignments (e.g., moving an entire motif to another position) exist.

**A2.** Short Gaps**.** Any genuine local indel has length strictly less than $l$.

**A3.** Gap Cost Threshold**.** For any possible set of local substitutions or indels, the maximal loss in score ($\Delta_{\max}$) caused by these local changes is strictly less than the cost of introducing a gap spanning an entire motif or multiple motifs. Formally, for any single motif, if the maximal score loss due to internal substitutions/indels is $\Delta_{\max}$, it must satisfy

$$\Delta_{\max} < -\gamma \cdot l.$$

Intuitively, it should not be favorable to "shift" an entire motif from one copy to another to increase alignment score.

**Theorem** (Equivalence of Segmented Alignment)

Under assumptions A1–A3: Consider $R$ as $c$ consecutive motif blocks. Perform precise local alignment only on blocks that differ from $t$ (i.e., copies with substitutions or indels). "Reuse/skip" the blocks that are identical to $t$. Then, the global alignment constructed by concatenating these local alignments: Is equivalent to a standard global alignment of $R$ and $Q$ in terms of total matches, mismatches, insertions, deletions, and overall alignment score, and Produces the same alignment path at block boundaries.

**Proof** (Sketch)

1) Each non-variant copy of $R$ is exactly equal to $t$. In a full global alignment, the alignment of this copy to the corresponding segment in $Q$ is locally optimal and deterministic.
2) If a copy $t$were misaligned to an adjacent copy, it would require a net gap of at least length $l$ (shifting the entire copy across the repeat unit), incurring a cost of at least $-\gamma l$. By assumption **A3**, this cost exceeds the maximal score loss from correcting local substitutions/short indels. Therefore, such shifts

cannot be globally optimal.

3) Consequently, in any globally optimal alignment, all unchanged copies are aligned in-place, and only blocks with local variations require detailed alignment.
4) This allows decomposition of the global problem into independent local alignment tasks: align each variant copy (length $\sim l$) individually, while treating unchanged copies as placeholders.
5) Concatenating the local alignments produces a global alignment whose path matches the full-length alignment at block boundaries, and whose counts of matches, mismatches, insertions, and deletions are identical to those obtained from a full-length global alignment.

**Note**.

Assumption **A3** is crucial: if gap penalties are too small ($|\gamma|$ very small), long gaps might be favorable compared to local substitutions, allowing cross-block shifts and invalidating the segmented alignment strategy. In practice, typical gap penalties and the biological cost of short indels are sufficient to satisfy **A3**.

The standard Needleman–Wunsch global alignment algorithm has a time complexity of $O(nm)$ for sequences of lengths $n$ and $m$ (assuming a basic dynamic programming implementation), and a space complexity of $O(nm)$ — which can be reduced to $O(n+m)$ using the Hirschberg algorithm. For a full-length alignment between the reference $R$ and the query $Q$, the computational cost is:

$$T_{\text{full}} = O(nm) = O\big((cl) \cdot m\big)$$

Under our segmented alignment strategy, assume that only a fraction $p$ (where $0 \le p \le 1$) of the motif copies in the query sequence $Q$ differ from those in the reference. That is, approximately $pc$ copies require precise local alignment. If these variant copies are concatenated into several segments, the total length of all segments is approximately: $L_{\text{var}} \approx (pc)l$. Then, the main computational cost of the segmented alignment can be approximated as:

$$T_{\text{seg}} = O(L_{\text{var}} \cdot m')$$

where $m'$ is the effective target length aligned against these variant segments (in the simplest case, $m'$ is roughly equal to $L_{\text{var}}$ or of the same order as the original $m$). Roughly comparing both cases, and assuming that both sides of the alignment are of similar length scale, we can approximate:

$$T_{\text{seg}} = O((pcl)^2) = p^2 \cdot O((cl)^2)$$

Thus, when $p \ll 1$, the theoretical speedup is about $1/p^2$ — that is, reducing the alignment length scale to a fraction $p$ of the original results in a quadratic reduction in computational cost. In practice, the actual acceleration is somewhat smaller than the theoretical value due to segment stitching, indexing overhead, and boundary processing, but the overall gain remains substantial when $p$ is small.

Even when we perform a standard global dynamic programming (DP) alignment for a given segment, the conventional procedure is to complete the DP matrix first and then perform traceback to count the numbers of matches, mismatches, insertions, and deletions. Below, we introduce a more direct method: by maintaining the necessary quantities during the bottom-up DP process (or by keeping both a scoring matrix and a "diagonal-step count" matrix), we can ultimately solve for $N_{\text{match}}, N_{\text{mismatch}}, N_{\text{ins}}, N_{\text{del}}$ algebraically — without traceback.

For two sequences $A$ (length $n$) and $B$ (length $m$), define the following quantities under the optimal global alignment:

1) $N_{\text{match}}$: number of matching bases (equal diagonal steps);
2) $N_{\text{mismatch}}$: number of mismatching bases (unequal diagonal steps);
3) $N_{\text{ins}}$: number of inserted bases (total number of vertical steps in the alignment);
4) $N_{\text{del}}$: number of deleted bases (total number of horizontal steps in the alignment).

Clearly, the following identities hold (counting the total steps along the alignment path):

$$\begin{cases} N_{\text{diag}} \equiv N_{\text{match}} + N_{\text{mismatch}} \\ N_{\text{diag}} + N_{\text{del}} = n, \\ N_{\text{diag}} + N_{\text{ins}} = m \end{cases}$$

(The first equation defines $N_{\text{diag}}$ as the total number of diagonal steps; the second and third equations follow from the path covering the full lengths of both sequences.) During DP computation, $N_{\text{diag}}$ can be directly obtained (explained below). Thus, we have four equations in total:

$$\begin{cases} N_{\text{match}} + N_{\text{mismatch}} = N_{\text{diag}}, \\ N_{\text{diag}} + N_{\text{del}} = n, \\ N_{\text{diag}} + N_{\text{ins}} = m, \\ \alpha N_{\text{match}} + \beta N_{\text{mismatch}} + \gamma(N_{\text{ins}} + N_{\text{del}}) = S_{\text{align}}, \end{cases}$$

where $S_{\text{align}}$ is the final alignment score.

From the first three equations, we can eliminate $N_{\text{ins}}, N_{\text{del}},$ and $N_{\text{mismatch}}$ to obtain a single linear equation in $N_{\text{match}}$:

$$\begin{cases} N_{\text{ins}} = m - N_{\text{diag}}, \\ N_{\text{del}} = n - N_{\text{diag}}, \\ N_{\text{mismatch}} = N_{\text{diag}} - N_{\text{match}}. \end{cases}$$

Substituting into the scoring equation gives:

$$S_{\text{align}} = \alpha N_{\text{match}} + \beta\left(N_{\text{diag}} - N_{\text{match}}\right) + \gamma\left(m + n - 2N_{\text{diag}}\right).$$

Simplifying, we have:

$$(\alpha - \beta)N_{\text{match}} = S_{\text{align}} - \beta N_{\text{diag}} - \gamma\left(m + n - 2N_{\text{diag}}\right).$$

Therefore, when $\alpha \neq \beta$:

$$N_{\text{match}} = \frac{S_{\text{align}} - \beta N_{\text{diag}} - \gamma\left(m + n - 2N_{\text{diag}}\right)}{\alpha - \beta}$$

Then, the remaining quantities can be obtained directly:

$$N_{\text{mismatch}} = N_{\text{diag}} - N_{\text{match}}, N_{\text{ins}} = m - N_{\text{diag}}, N_{\text{del}} = n - N_{\text{diag}}.$$

Hence, as long as we know $S_{\text{align}}$, $n$, $m$, and $N_{\text{diag}}$, we can analytically determine all alignment statistics in a single pass — without performing traceback.

During the standard dynamic programming (DP) procedure (filling the scoring matrix $S[i,j]$), we can simultaneously maintain an auxiliary matrix $D[i,j]$ of the same size, where $D[i,j]$ represents the number of diagonal steps (diagonal count) in the optimal path from the origin $(0,0)$ to $(i,j)$ that corresponds to $S[i,j]$.

1) Initialization

$$S[0,0] = 0, D[0,0] = 0$$

$$S[i,0] = i\gamma, D[i,0] = 0$$

$$S[0,j] = j\gamma, D[0,j] = 0$$

**2)** Recurrence Relations

When filling in $S[i,j]$, consider the three possible sources:

$$\begin{cases} \text{Diagonal:} & s_{\text{diag}} = S[i-1,j-1] + \text{score}(A_i, B_j), \\ \text{Up (deletion):} & s_{\text{up}} = S[i-1,j] + \gamma, \\ \text{Left (insertion):} & s_{\text{left}} = S[i,j-1] + \gamma. \end{cases}$$

Then,

$$S[i,j] = \max\{s_{\text{diag}}, s_{\text{up}}, s_{\text{left}}\}.$$

At the same time, we update $D[i,j]$ according to the source that produced the optimal score $S[i,j]$: (if multiple sources tie, a deterministic priority rule should be used to ensure reproducibility.)

$$D[i,j] = \begin{cases} D[i-1,j-1] + 1, & \text{if } S[i,j] = s_{\text{diag}}, \\ D[i-1,j], & \text{if } S[i,j] = s_{\text{up}}, \\ D[i,j-1], & \text{if } S[i,j] = s_{\text{left}}. \end{cases}$$

(When $S[i,j] = s_{\text{diag}}$, one can also check whether $A_i = B_j$ to distinguish matches from mismatches; however, for computing $N_{\text{diag}}$, we only need the total number of diagonal steps.)

3) Complexity and Output

This auxiliary update requires only constant time per cell (a few comparisons and assignments), so the total time complexity remains $O(nm)$, and if we store the full matrices, the space complexity is also $O(nm)$.

After completing the DP process, we simply take:

$$N_{\text{diag}} = D[n,m], S_{\text{align}} = S[n,m],$$

and substitute these values into the algebraic formulas derived earlier to obtain all alignment statistics — without performing traceback.

### Supplementary Notes

#### Supplementary Note 1: Read Assembly and Assembly Evaluation of the Lung Cancer Sample

We used whole-genome sequencing data from a lung adenocarcinoma sample generated by the Medical Genome Sciences Laboratory, University of Tokyo (UT-MGS) (see Supplementary Table 1). This tumor sample was assembled *de novo*, and the assembly results were further evaluated and compared across 82 publicly available human genomes from the 1000 Genomes Project and the Human Pangenome Project.

First, the raw sequencing reads were quality-controlled using Chopper (version 0.9.2) with the following command:

```
chopper -q 7 -l 5000 -t 32 -i <input.fastq>
```

After quality control, the genome assembly was performed using Flye (version 2.9.5-b1801) with the command:

```
flye --nano-raw --genome-size 3.2g --threads 32 -i <input.fastq> -o <output_dir>
```

The resulting draft assembly was further polished using Medaka (versuib 2.0.1) to correct base-level errors:

```
medaka_consensus -o consensus -t 32 -b 50 -m r941_prom_hac_g507-i -d <draft.fa>
```

To rapidly assess the completeness of the lung cancer genome assembly, we employed BUSCO (version 5.8.2) with the primates_odb12 database (see Supplementary Table 3) using the following command:

```
busco -o busco -c 32 -l primates_odb12 -m genome --offline -i <assembly.fa>
```

Supplementary Table 4 BUSCO-based evaluation of genome assembly completeness in lung cancer samples

| Sample | C(%) | S(%) | D(%) | F(%) | M(%) |
|---|---|---|---|---|---|
| DRR171429 | 2.2 | 2.2 | 0 | 0.1 | 97.6 |
| DRR171430 | 14.0 | 13.9 | 0.1 | 0.5 | 85.5 |
| DRR171431 | 90.6 | 89.0 | 1.6 | 0.7 | 8.6 |

| | | | | | |
|---|---|---|---|---|---|
| DRR171432 | 82.2 | 80.6 | 1.6 | 0.9 | 16.9 |
| DRR171433 | 23.4 | 23.2 | 0.2 | 1.2 | 75.4 |
| DRR171452 | 58.9 | 57.5 | 1.4 | 1.6 | 39.6 |
| DRR171453 | 96.9 | 95.8 | 1.1 | 0.3 | 2.7 |
| DRR171454 | 78.6 | 77.3 | 1.3 | 1.2 | 20.2 |
| DRR203145 | 99.6 | 98.8 | 0.8 | 0.2 | 0.2 |

Complete BUSCOs (C) refers to the proportion of full-length BUSCO genes identified; Single-copy BUSCOs (S) are complete genes found in single copy; Duplicated BUSCOs (D) are complete genes found in multiple copies; Fragmented BUSCOs (F) are partial gene fragments; Missing BUSCOs (M) are BUSCO genes not identified in the assembly. n indicates the total number of BUSCO groups searched using the *primates_odb12* dataset (n = 11,834).

To comprehensively evaluate the assembly quality of the lung cancer sample and other human individual samples, we performed standardized quality assessment on all 91 assemblies (including the 9 lung cancer samples and 82 publicly available human samples) using QUAST (version 5.3.0) (Supplementary Table 4), with T2T-CHM13v2.0 as the reference genome. The command used was:

```
quast.py <assemblies> -r GCA_009914755.4_T2T-CHM13v2.0_genomic.fna \
--large --threads 16 --eukaryote --fragmented -o <output_dir>
```

Compared to normal samples derived from healthy donors with deep sequencing coverage, the assemblies of lung cancer cell line samples were generally more fragmented, with lower coverage and poorer continuity. Nevertheless, several top-performing lung cancer samples (e.g., DRR171453, DRR203145) achieved nearly 90% genome fraction and megabase-level N50, indicating that with optimized sequencing and assembly parameters, tumor samples can still yield high-quality genome assemblies suitable for downstream structural variation analysis.

**Supplementary Note 2: Comprehensive Analysis of STR Quality Distribution**

To comprehensively characterize the quality distribution of true STRs and accurately determine the optimal quality threshold for STR identification in FastSTR, we obtained all STRs detected by TRF (alignment parameters: *2 5 7*) on the human reference genome T2T-CHM13. For each motif length from 1 to 8 bp, we randomly sampled 5,000–9,000 STRs as the test set.

We used the percentage of matches ($P_{match}$) and the percentage of insertions/deletions ($P_{indel}$) (defined below) to quantify STR quality.

Overall, as $P_{indel}$ increases and $P_{match}$ decreases, the distribution of tandem repeats rapidly converges toward 0. Approximately x% of STRs have $P_{indel}$ < 0.15, and X% have $P_{indel}$ > 0.80. Although the quality distributions vary across different motifs, the majority converge near $P_{indel}$ = 0.15 and $P_{match}$ = 0.80 **(Supplementary Fig.2,3)**. When repeating the same analysis using TRF parameters 2 3 5, the results remained nearly unchanged, indicating that alignment parameters have minimal impact on STR quality distribution. Using a composite quality metric $\rho = 1 - P_{indel} + P_{match}$ to evaluate STR distribution, we found that different motifs exhibit distinct quality distributions and minimum values (ranging from a maximum of 1.80 to a minimum of 1.72) **(Supplementary Fig.4)**. However, all were significantly higher than the theoretical lower bound (1 − 0.15 + 0.80 = 1.65), suggesting that under the given alignment constraints, low-quality STRs with both high indel rates and low match rates do not occur.

**Supplementary Note 3: Validation of STR Identification Strategy and Parameter Settings in FastSTR**

1) Density-based clustering for fuzzy repeat region detection

The outcome of density-based clustering depends critically on parameter selection. Based on the observed STR quality distribution and the definition of anchor points, we empirically estimated the optimal parameters. Considering that FastSTR identifies STRs with a minimum length of 25 bp and that terminal regions may contain noise, both the clustering radius and minimum sample size were set to 14.

Tests showed that clustering performance remained stable for parameter values near 14**(Supplementary Fig.5)**.

In the test dataset, most STRs with flanking regions shorter than 250 bp were successfully clustered. Therefore, during STR identification, we extended each candidate STR by 500 bp on both sides and re-evaluated the extended regions to detect potential adjacent STRs that may have been erroneously merged due to close proximity.

2) Handling interrupted clustering and merging of adjacent STRs

We further observed that certain STRs were fragmented during clustering due to low-quality sequence segments. To assess this effect, we applied density-based clustering to all STRs identified by TRF on the T2T reference genome. The results indicated that interrupted STRs accounted for less than 0.5% of each chromosome**(Supplementary Fig.6 a)**, with slightly higher proportions observed only on chromosomes containing a large number of long, low-quality STRs.

Length distribution analysis of the interrupted segments revealed that 99% were shorter than 100 bp**(Supplementary Fig.6 a)**, with most concentrated below 50 bp. Accordingly, in the STR merging phase, FastSTR considers adjacent STRs separated by less than 100 bp as potential candidates for merging into a single continuous STR, ensuring accurate reconstruction of the full repeat region.

3) Filtering false-positive repeat regions using leading-base density

We determined motif-length–specific thresholds for leading-base density based on 97% and 99% confidence intervals derived from the test dataset. To assess their validity, 5,000 simulated STRs with motif lengths from 1 bp to 8 bp were generated and subjected to chi-square testing. Results showed that over 98% of true STRs had leading-base densities above the threshold, whereas more than 98% of false STRs fell below it**(Supplementary Fig.7 a,b)**. These findings indicate that the selected thresholds effectively distinguish genuine STRs from false positives, substantially reducing spurious detections.

4) Accurate motif recognition using the N-Gram model and Markov chain

We extracted tandem repeats (TRs) with motif lengths ranging from 1 bp to 30 bp identified by TRF from the human T2T reference genome, selecting at least 5,000 TRs per motif length for analysis. Using a pure N-Gram model (without the Markov component), we predicted representative motifs and compared them with TRF annotations. The results revealed that STRs with motifs ≤ 8 bp achieved recognition accuracies above 80%, whereas those with motifs > 9 bp showed a sharp decline**(Supplementary Fig.8 a)**. This suggests that the N-Gram–based method is particularly suited for canonical STRs. When the N-Gram model was combined with a Markov chain for motifs of 5–8 bp, the recognition accuracy improved substantially**(Supplementary Fig.8 b)**, approaching the performance observed for 1–4 bp motifs. These results demonstrate that the Markov chain captures intrinsic structural dependencies within motifs, thereby enhancing motif recognition accuracy.

5) Efficient alignment of ultra-long STRs using a segmented global alignment algorithm

To evaluate the computational efficiency of the segmented global alignment algorithm, we simulated 3,000 ultra-long STRs ranging from 7 kb to 70 kb and compared results with those from a conventional local alignment algorithm. Alignment accuracy was assessed using the absolute sum of indel and match percentage deviations (lower values indicating higher accuracy). On average, the segmented algorithm reduced runtime by 58.694 s, with 89.6% of alignments saving more than 5 s**(Supplementary Fig.9 a)**. Additionally, 99.57% of alignments exhibited a combined error below 0.06**(Supplementary Fig.9 b)**. These results confirm that the segmented global alignment algorithm achieves substantial computational acceleration while maintaining near-identical alignment precision for ultra-long STRs.

**Supplementary Note 4: Systematic Comparison of STR Detection Tools**

To comprehensively evaluate the performance of FastSTR, we conducted a systematic comparison with three widely used STR detection tools — TRF (version 4.09.1), mreps (version 2.6), T-reks (version 1.3), and TRASH (version 1.2).

Because mreps and T-reks lack parallel processing capabilities and are relatively slow, each genome was split by chromosome, and multiple instances were run in parallel. The total runtime was estimated by summing the execution times of all subprocesses, and the peak memory usage was taken as the maximum among them. This approximation closely reflects realistic performance. Runtime and memory usage were measured using the /usr/bin/time command, while peak memory for TRASH (which supports multithreading) was assessed using smem. Commands used for each tool:

```
trf fasta 2 5 7 80 10 50 2000 -d -h -l 10
```

```
faststr 2 5 7 3 15 80 50 fasta -p 72
```

```
mreps -res 5 -exp 2.0 -minsize 15 -maxp 8 -allowsmall -fasta fasta
```

```
java -Djava.awt.headless=true -jar T-Reks.jar -msaMode=x -muscle=muscle -
clustal=clustalw -infile=fasta
```

```
TRASH fasta --o outputdir --par 70 arguments
```

Using STRs detected by TRF as the ground truth, we examined the distribution of right-boundary deviations for each tool. The deviation pattern was symmetric with respect to the left boundary**(Supplementary Fig.10)**, and most STR boundaries identified by FastSTR were perfectly aligned (0 bp deviation), indicating superior boundary precision.

When comparing copy numbers between FastSTR and TRF across 13 representative species, the majority of STRs showed differences within ±1, while approximately 5% differed by 1–2 copies, and 5–10% differed by more than 5 copies**(Supplementary Fig.11)**. Notably, STRs with a 7 bp motif in *Homo sapiens* and 4 bp motifs in *Zea mays* exhibited larger discrepancies, likely due to ambiguous repeat boundaries that different algorithms interpret differently.

We further examined novel STRs uniquely identified by FastSTR, including (i) overlapping regions missed by TRF for motifs < 8 bp, and (ii) non-overlapping regions representing entirely new STR loci. Validation was performed using the T2T STR reference annotation**(Supplementary Fig.12 a)**, as well as T-reks and mreps tools. The results showed that in known overlapping regions, 36,861 out of 38,957 STRs were validated, yielding a validation rate of 94.62%; in novel overlapping regions**(Supplementary Fig.12 b)**, 3,629 out of 3,642 STRs were confirmed, corresponding to a validation rate of 99.64%. Collectively, these results demonstrate that FastSTR not only achieves superior computational efficiency but also exhibits high reliability in identifying both known and novel STRs across species.

**Supplementary Note 5 : Complementary Analysis of STR Entries Between FastSTR, TRF, and the T2T STR Reference Annotation**

We merged STR regions identified by FastSTR, the T2T STR reference annotation, and TRF into a set of non-overlapping genomic intervals using the following command:

```
bedtools merge -i str.bed > str.merged.bed
```

Since the bedtools intersect function determines overlap counts based on the file specified with the -a parameter, the number of overlapping intervals may vary depending on input order. To ensure consistency, we computed all pairwise overlap counts across different input orders and selected the minimum value as the effective overlap count between two sources (A and B):

```
bedtools intersect -u -a Astr.merged.bed -b B.merged.bed > AandBstr.merged.bed
```

After obtaining pairwise overlaps, we extended the same principle to compute the minimum shared overlaps among all three annotation sources, representing the commonly identified STR loci.

A Venn diagram of the three datasets revealed that the T2T STR reference contained the largest number of unique STRs (305,031 entries) **(Supplementary Fig.13 a)**. However, subsequent local alignment analysis showed that 99.85% of these unique STRs had alignment scores below 50**(Supplementary Fig.13 b)**. Given that both FastSTR and TRF require a minimum alignment score of 50 to qualify as valid STRs, these entries were excluded from further consideration. This explains the apparent excess of unique STRs identified only by the T2T STR reference.

We next examined the genome-wide distribution of STRs jointly identified by all three methods **(Supplementary Fig.14)**. STR density was markedly lower near centromeric and telomeric regions, while other genomic regions showed relatively uniform distributions. Notably, the Y chromosome exhibited a significantly higher STR density compared to other chromosomes.

For STRs uniquely identified by FastSTR but validated by the T2T STR reference, we analyzed the motif composition. The predominant category was HSATII**(Supplementary Fig.15)**, followed by a set of complex tandem repeats composed of shorter repeating subunits, a pattern that TRF often fails to resolve effectively.

Finally, we compared the sequence quality (insertion/deletion and match percentages) of STRs identified by FastSTR and TRF, stratified by whether they were validated by the T2T STR reference. The results, visualized as hexbin plots**(Supplementary Fig.16,17)**, show that STRs identified by FastSTR exhibit a more compact and consistent quality distribution, particularly those corresponding to HSATII regions, underscoring FastSTR's robustness in identifying complex repeat structures.

**Supplementary Note 6 : Population-scale Analysis of STR Variation Using FastSTR**

We first applied FastSTR to identify STRs across 82 high-quality genome assemblies. Each assembly was then aligned to both the T2T and GRCh38 (HG38) reference genomes using minimap2 with the following parameters:

```
minimap2 -cx asm20 --cs -m 10000 -z 10000,50 -r 50000,2000000 -t 30 --end-bonus=100 --rmq=yes -O 5,56 -E 4,1 T2T.fna asm.fa > asm.t2t.paf
```

```
sort -k6,1 -k8,2n asm.t2t.paf > asm.t2t.srt.paf
```

```
minimap2 -cx asm20 --cs -m 10000 -z 10000,50 -r 50000,2000000 -t 30 --end-bonus=100 --rmq=yes -O 5,56 -E 4,1 GRCh38.fasta asm.fa > asm.hg38.paf
```

```
sort -k6,1 -k8,2n asm.hg38.paf > asm.hg38.srt.paf
```

Next, we lifted over STR coordinates identified by FastSTR to the T2T and HG38 reference genomes using the liftover function in paftools.js:

```
paftools.js liftover -l20 asm.t2t.srt.paf str.bed > t2t.liftover.bed
```

```
paftools.js liftover -l20 asm.hg38.srt.paf str.bed > hg38.liftover.bed
```

Structural variants relative to the two references were detected using paftools.js call, retaining only variants with quality ≥ 20 and containing indels:

```
paftools.js call -asm.t2t.srt.paf -f T2T.fna -q 60 -L 10000 > asm.t2t.var.txt
```

```
paftools.js call -asm.hg38.srt.paf -f GRCh38.fasta -q 60 -L 10000 > asm.hg38.var.txt
```

We calculated Jaccard similarity of STR variations within and between populations, based on both all detected variants and novel STR variants uniquely identified by FastSTR **(Supplementary Fig.18 a,b)**. The intra-population similarity was consistently higher than inter-population similarity. Populations from the Americas exhibited higher similarity with East Asian, African, and European groups, whereas African populations showed lower similarity with East, South Asian, and European groups. These results align with established models of human migration and population differentiation, reaffirming the genetic distinctiveness of African lineages and the historical admixture of American populations.

Finally, we analyzed FastSTR-specific STR variants across family trios and among populations, quantifying super-population-specific and individual-specific STR variants**(Supplementary Fig.19,20)**. The Jaccard similarity of STR variation within families was substantially higher than between unrelated families. Together, these findings demonstrate that FastSTR provides high-resolution, population-scale insights into STR diversity, supporting both evolutionary and medical genomic applications.

**Supplementary Note 7: Detection of Pathogenic STR Variants in Lung Cancer Using FastSTR**

We first applied FastSTR to identify STR sequences in nine cancer samples, including eight lung cancer and one breast cancer sample. STR intervals from all cancer samples were then merged using bedtools to obtain a set of non-overlapping STR regions. Following the same procedure as in the assembly-based analysis, we aligned each cancer genome to the T2T reference genome (CHM13v2.0) using minimap2, and mapped STR interval positions to the reference using paftools, which also detected sequence variants between cancer samples and the T2T genome.

To assess the potential functional impact of these variants at the amino acid level, we performed annotation with ANNOVAR. To ensure compatibility with the latest T2T reference, we used the T2T transcript annotation file hs1_refGene.txt.gz and executed the following command:

```
perl table_annovar.pl str_variants.avinput /annovar/humandb/ -buildver hs1 -out
str_annotation -protocol refGene -operation g -nastring . -polish
```

We further conducted GO enrichment analyses for genes containing STR variants located within coding sequences (CDS) and untranslated regions (UTRs) using g:Profiler. Terms were filtered to include between 10 and 500 genes, and only those with adjusted $p$ values < 0.05 were retained. The results revealed significant enrichment of these genes in the biological process (BP), molecular function (MF), and cellular component (CC) categories, suggesting that STR variations may contribute to tumorigenesis through multiple functional pathways**(Supplementary Fig.21,22)**.

### Supplementary Figures

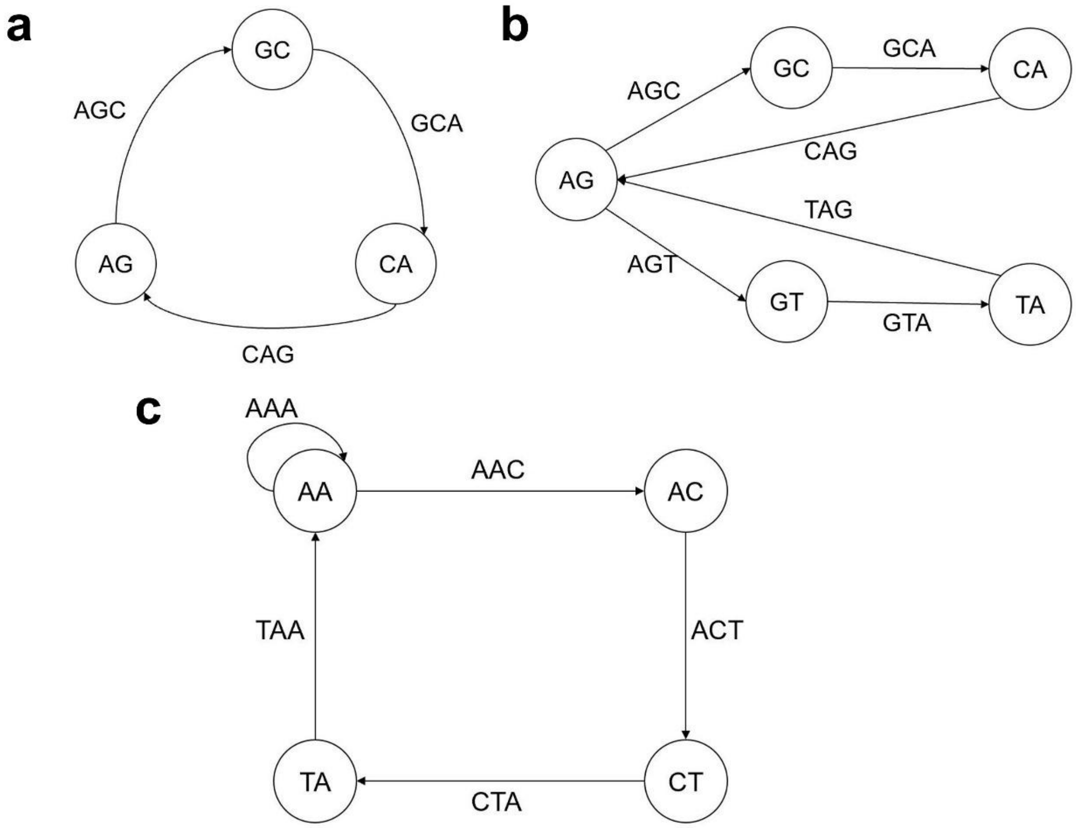


Supplementary Fig.1 a. The directed graph corresponding to the motif AGCAG is a trivial cycle. b. The motifs "AGCAGTAG" and "AGTAGCAG" correspond to the same directed graph, which contains a branching path. c. The motifs "AAACCCAA" and "CCCAAACC" correspond to the same directed graph, which contains two self-loop paths.

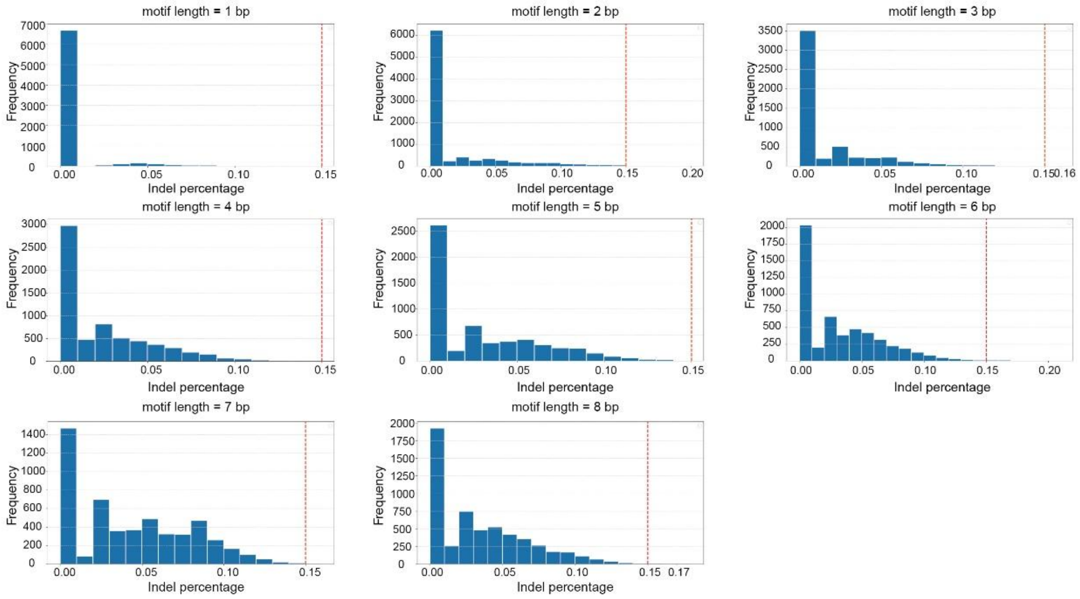


Supplementary Fig. 2 The distribution of indel percentages for STRs with motif lengths ranging from 1 to 8 bp. The statistics were calculated separately for each motif length, and the indel percentage of STRs for every motif length is less than 0.15.

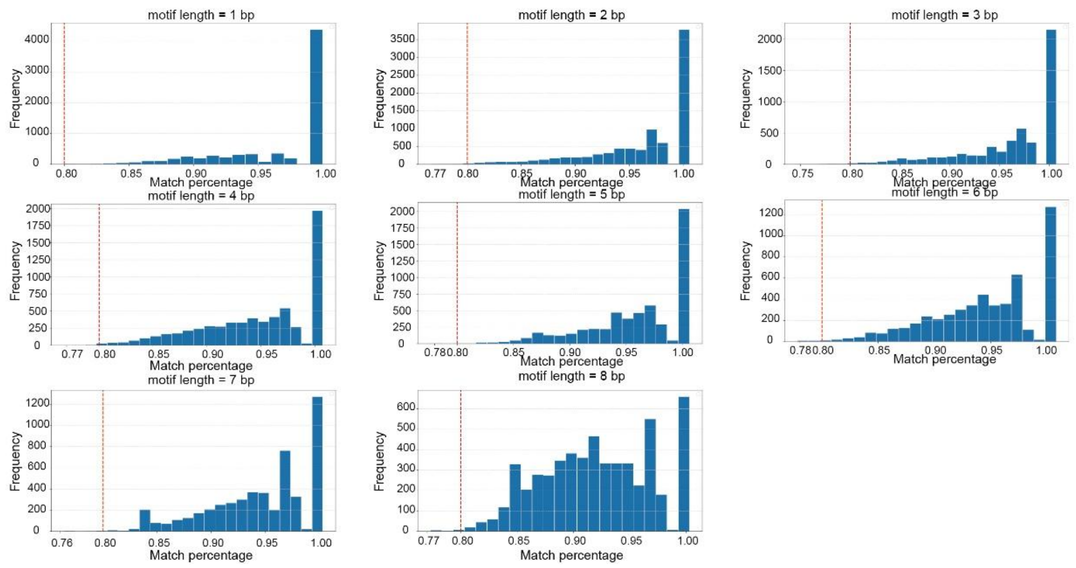


Supplementary Fig. 3 The distribution of match percentages for STRs with motif lengths of 1–8 bp. The analysis was performed for each motif length, and the match percentage of STRs in all cases exceeds 0.8.

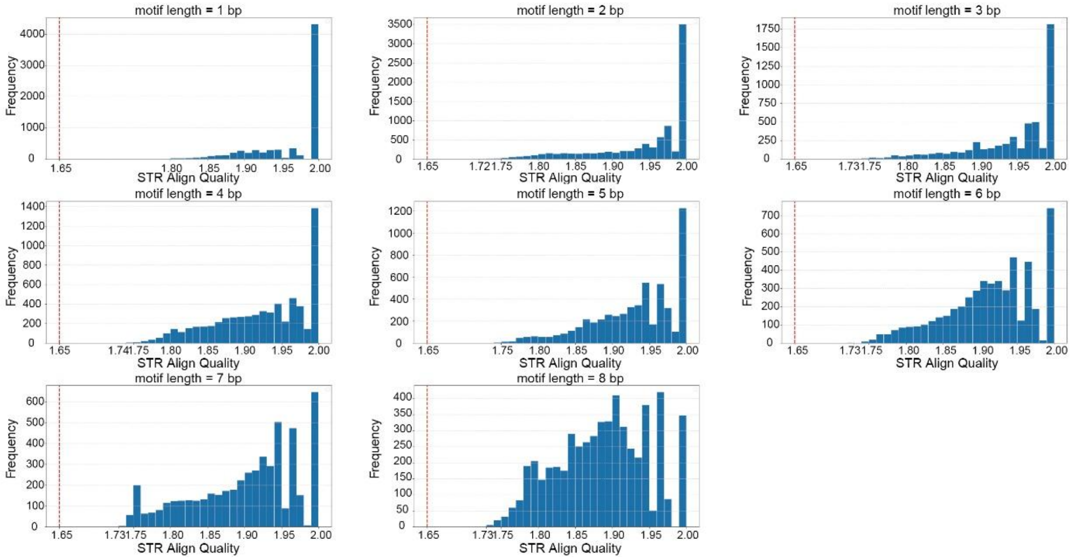


Supplementary Fig. 4 The quality distribution of STRs with motif lengths of 1–8 bp. The quality value was derived from both the indel and match percentages and was analyzed across different motif lengths. The results indicate that the quality of STRs for all motif lengths is substantially higher than 1.65, and the quality distributions vary among different motif lengths.

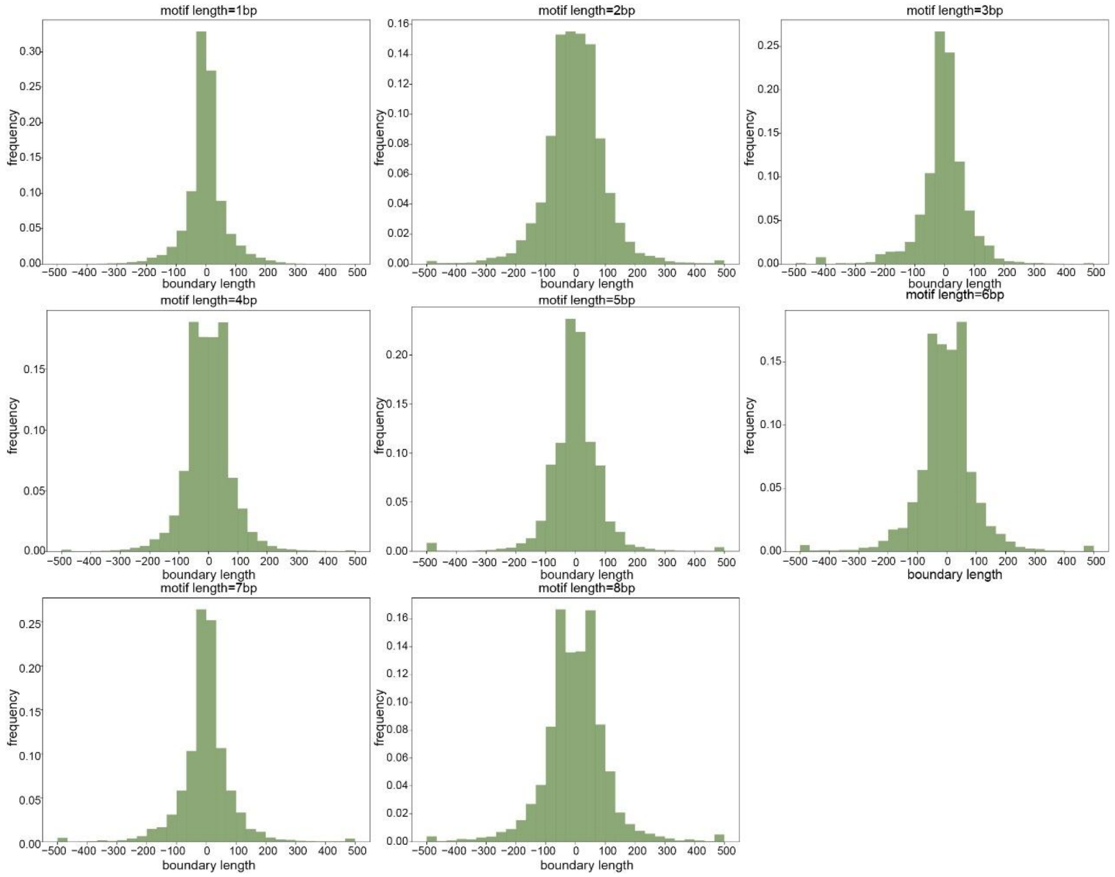


Supplementary Fig. 5 Length distribution of flanking sequences after density-based clustering for STRs with motif lengths ranging from 1 to 8 bp, including 500 bp of flanking regions on both sides. The statistics were computed separately for each motif length. Most STR flanking sequences are markedly enriched within 200 bp after density clustering.

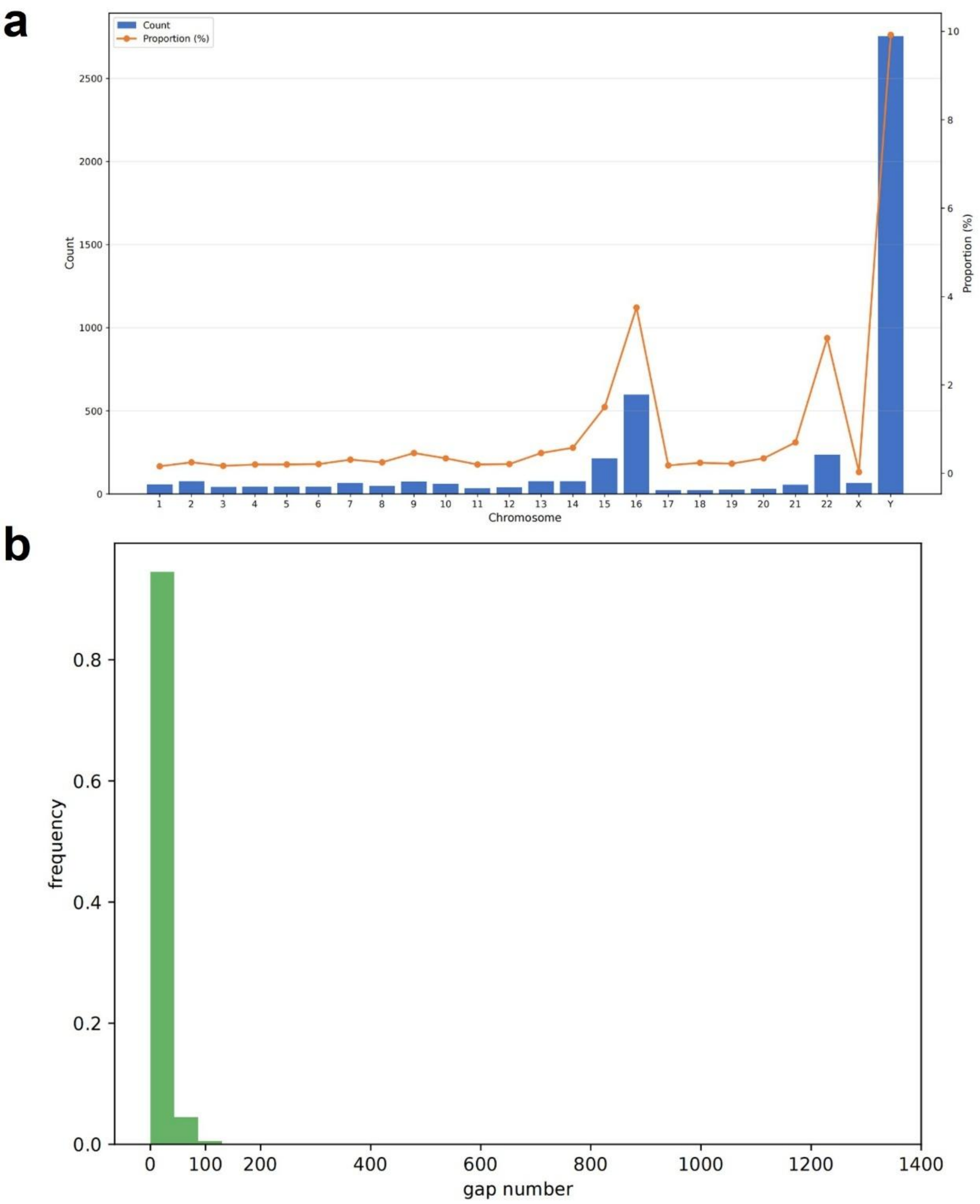


Supplementary Fig. 6 a. Proportion of disrupted STRs across chromosomes after density-based clustering of all STRs identified by TRF in the T2T reference genome. In most chromosomes, the proportion of disrupted STRs is less than 0.5%. b. Distribution of gap lengths among disrupted STRs. The majority of gap lengths are markedly enriched within 100 bp.

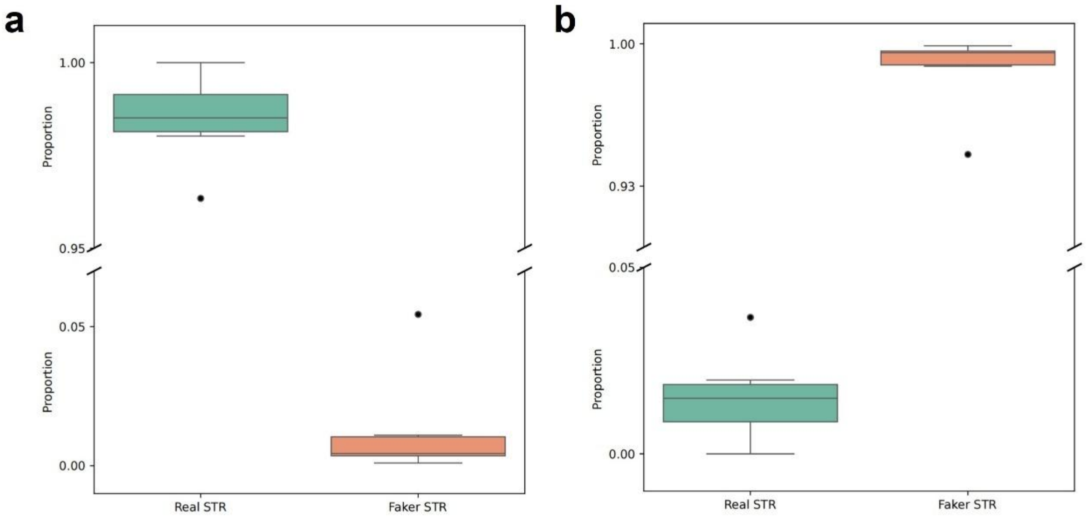


Supplementary Fig. 7 a. P Proportion of true and false STRs with motif lengths ranging from 1 to 8 bp that exceed the leading-base density threshold in randomly simulated datasets. Most true STRs are above the threshold, whereas almost no false STRs exceed it. b. In contrast to panel a, the proportions of STRs below the leading-base density threshold are shown.

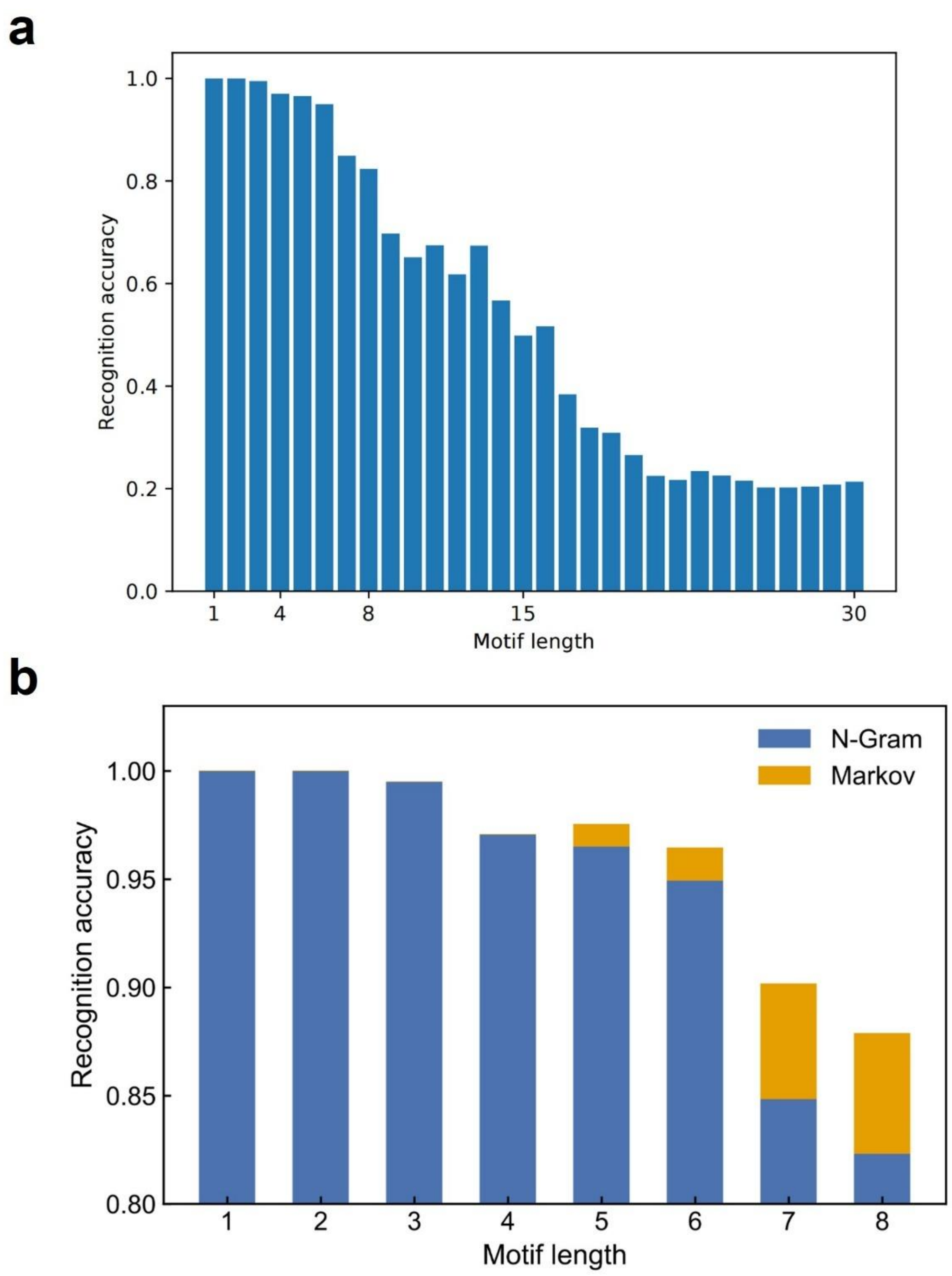


Supplementary Fig. 8 a. Motif identification performance of the N-Gram model across TRs with motif lengths ranging from 1 to 30 bp. The recognition accuracy markedly decreases when the motif length exceeds 8 bp, indicating that the N-Gram model is suitable only for motifs no longer than 8 bp. b. Improvement in motif identification accuracy for STRs with motif lengths of 5–8 bp after incorporating a Markov chain model. The enhancement is particularly pronounced for motifs of 7 bp and 8 bp.

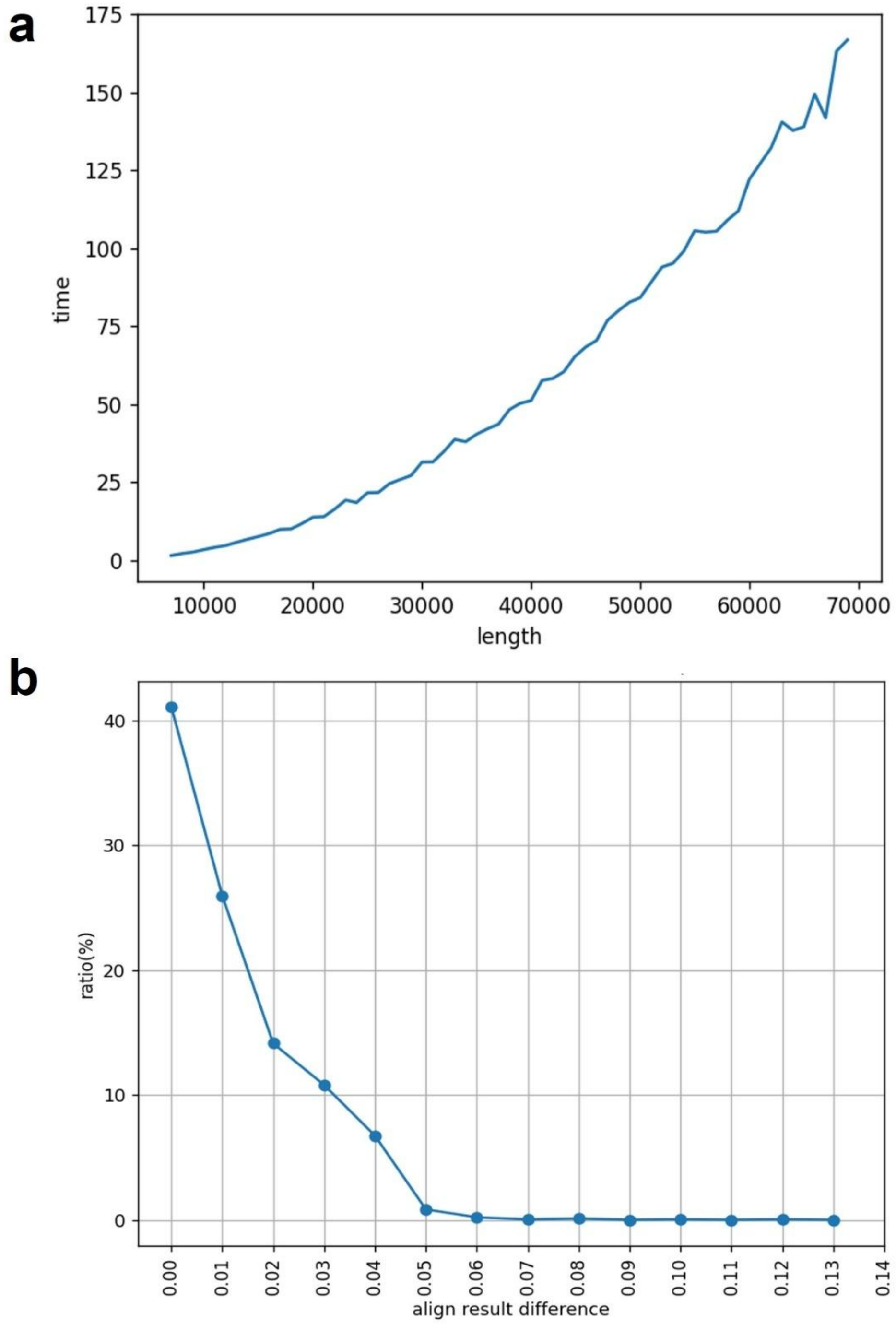


Supplementary Fig. 9 a. Performance evaluation of the segmented global sequence alignment algorithm compared with the conventional alignment algorithm using randomly simulated STRs ranging from 7,000 bp to 70,000 bp in length. b. Comparison of indel and match percentages identified by the segmented global alignment algorithm and the conventional alignment algorithm, showing similar detection results between the two methods.

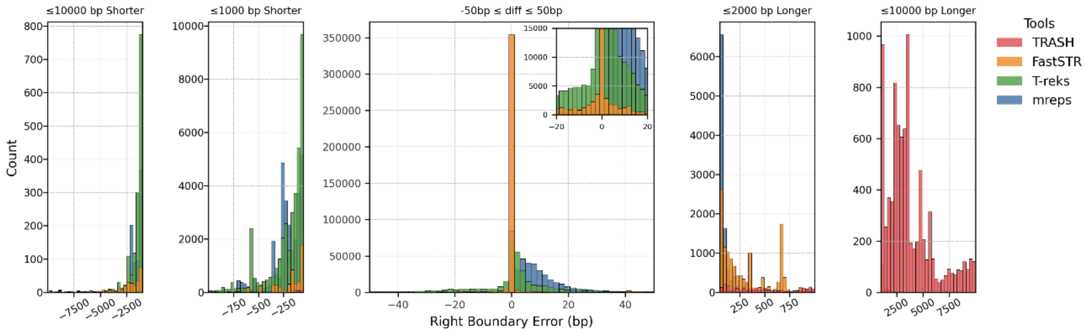


Supplementary Fig. 10 Distribution of right-boundary deviations of STRs identified by different tools relative to TRF in the human genome. FastSTR shows the smallest boundary deviation, with most differences concentrated around 0 bp.

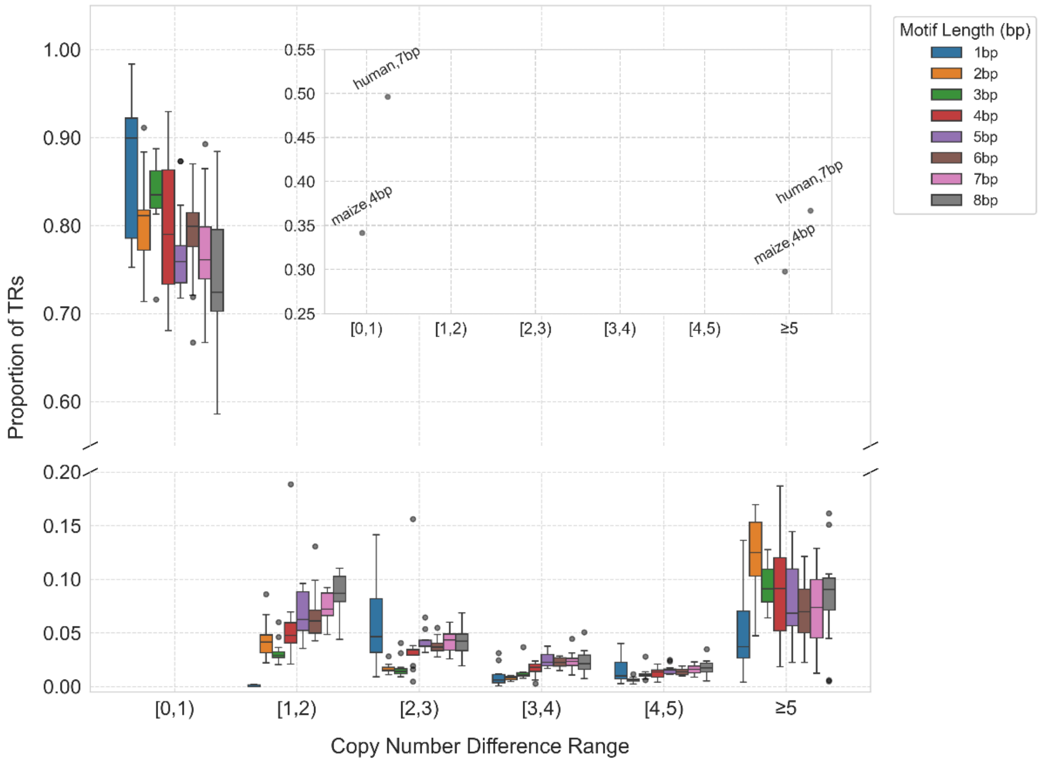


Supplementary Fig. 11 Distribution of copy number differences between STRs identified by FastSTR and TRF across 13 species, stratified by motif length. Most differences are concentrated within 0–1 bp, indicating high concordance between FastSTR and TRF. However, the 7 bp motifs in human and 4 bp motifs in maize exhibit larger discrepancies, reflecting lower consistency in copy number identification for these cases.

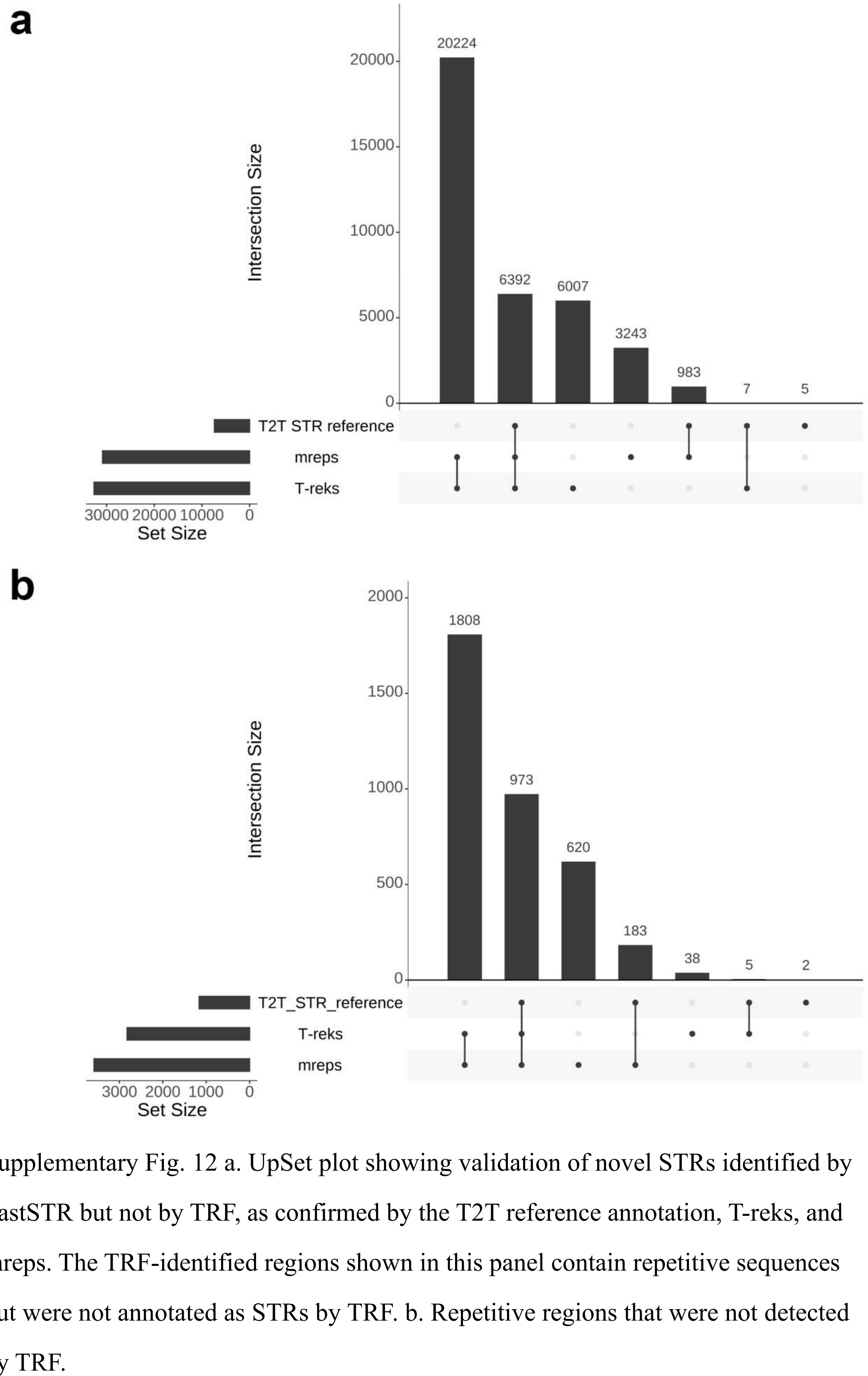


Supplementary Fig. 12 a. UpSet plot showing validation of novel STRs identified by FastSTR but not by TRF, as confirmed by the T2T reference annotation, T-reks, and mreps. The TRF-identified regions shown in this panel contain repetitive sequences but were not annotated as STRs by TRF. b. Repetitive regions that were not detected by TRF.

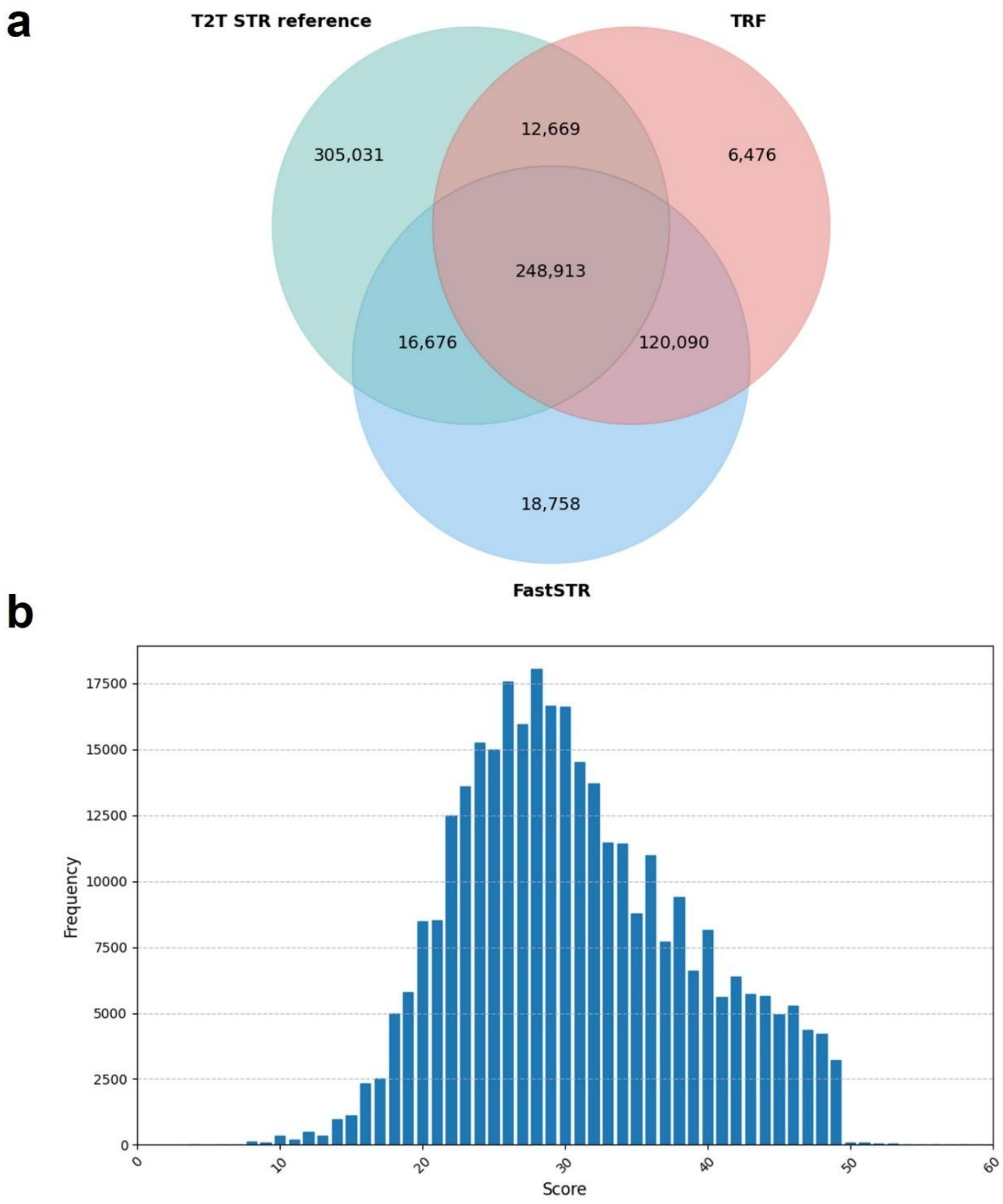


Supplementary Fig. 13 a. Venn diagram showing the number of overlapping STRs among FastSTR, the T2T reference STR annotation, and TRF-identified STRs. b. Distribution of alignment scores for STRs uniquely identified by the T2T reference annotation. Most of these STRs have alignment scores below 50, whereas STRs identified by TRF and FastSTR consistently exhibit alignment scores of 50 or higher.

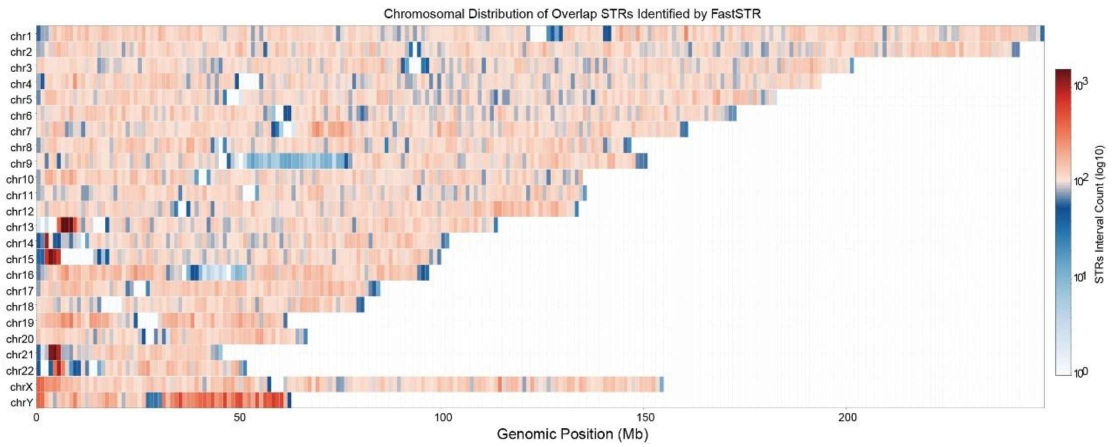


Supplementary Fig. 14 Genomic distribution of STRs annotated by FastSTR, the T2T reference STR annotation, and TRF across the human genome. STRs are less frequent near centromeric and telomeric regions, whereas a higher density is observed on the Y chromosome.

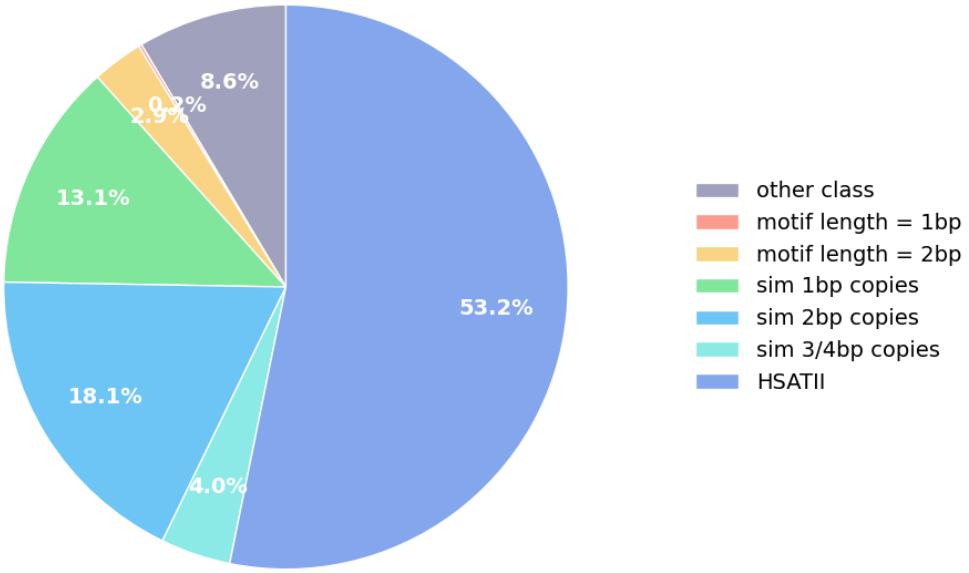


Supplementary Fig. 15 Distribution of motif structures for STRs uniquely identified by FastSTR relative to TRF and validated by the T2T reference STR annotation. Most motifs belong to the HASTII type or consist of tandemly repeated shorter repeat units forming similar structures.

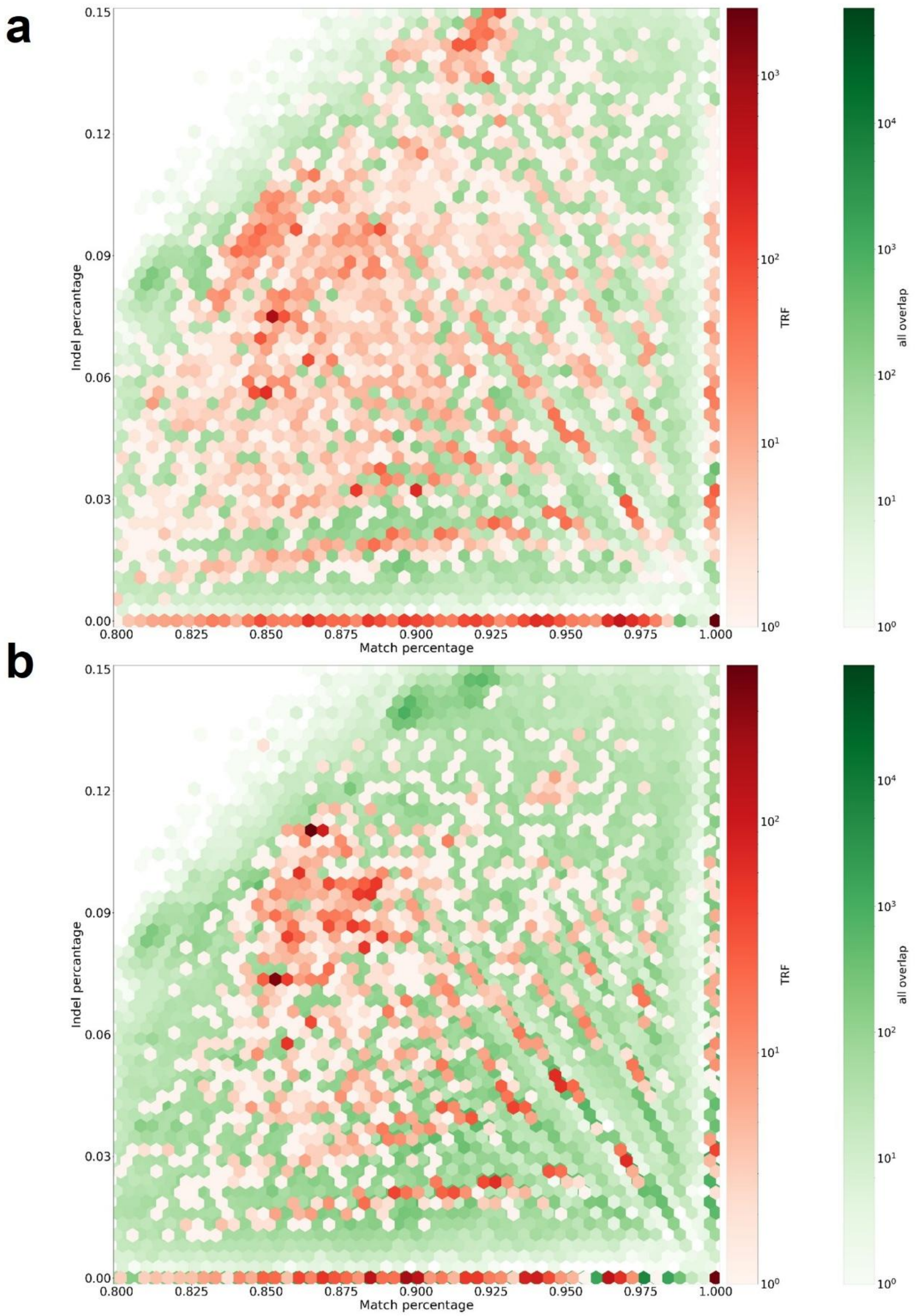


Supplementary Fig. 16 a. Distribution of quality scores for STRs identified by TRF but not by FastSTR, validated by the T2T STR reference annotation. b. Distribution of quality scores for STRs uniquely identified by TRF.

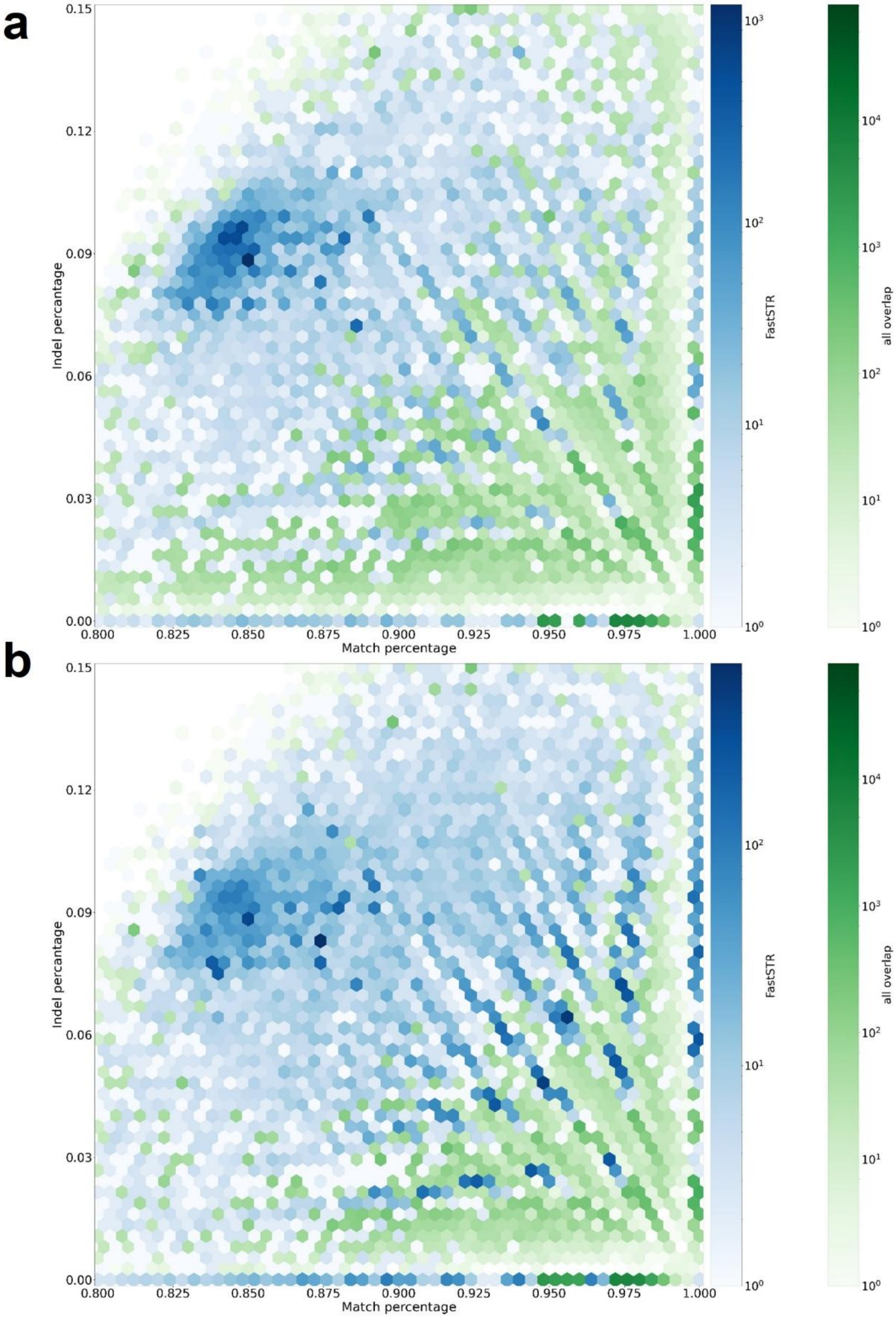


Supplementary Fig. 17 a. Distribution of quality scores for STRs identified by FastSTR but not by TRF, validated by the T2T STR reference annotation. b. Distribution of quality scores for STRs uniquely identified by FastSTR.

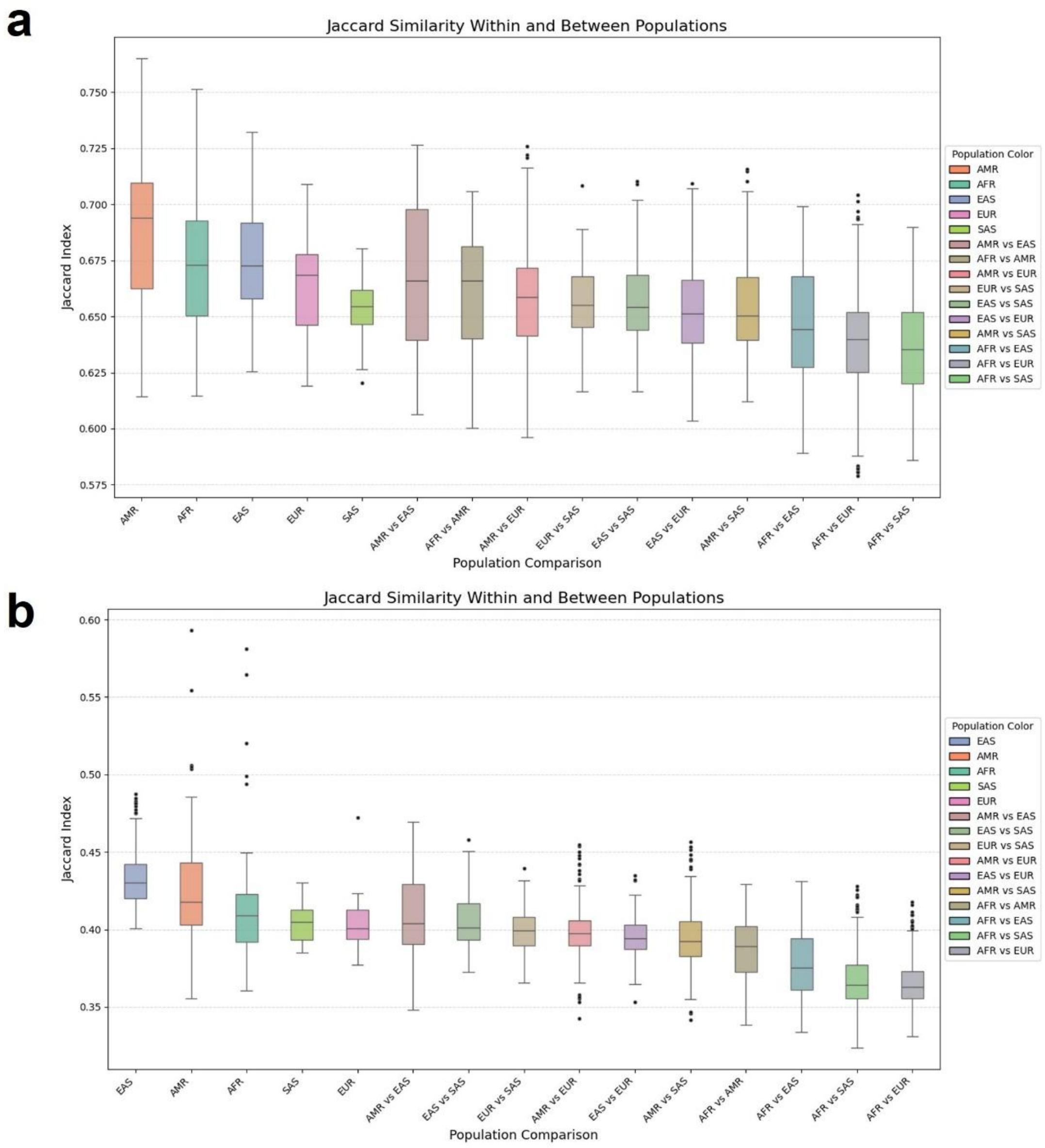


Supplementary Fig. 18 a. Boxplots of pairwise Jaccard similarities of all STR variants between individuals from different populations. Individuals within the same population show higher similarity than those between populations. Notably, the similarity between the American population and East Asian, European, and African populations is relatively high, whereas African individuals show the lowest similarity with East Asian, South Asian, and European populations. b. Boxplots of pairwise Jaccard similarities of novel STR variants uniquely identified by FastSTR across populations. The pattern of similarity is consistent with that observed for all STR variants.

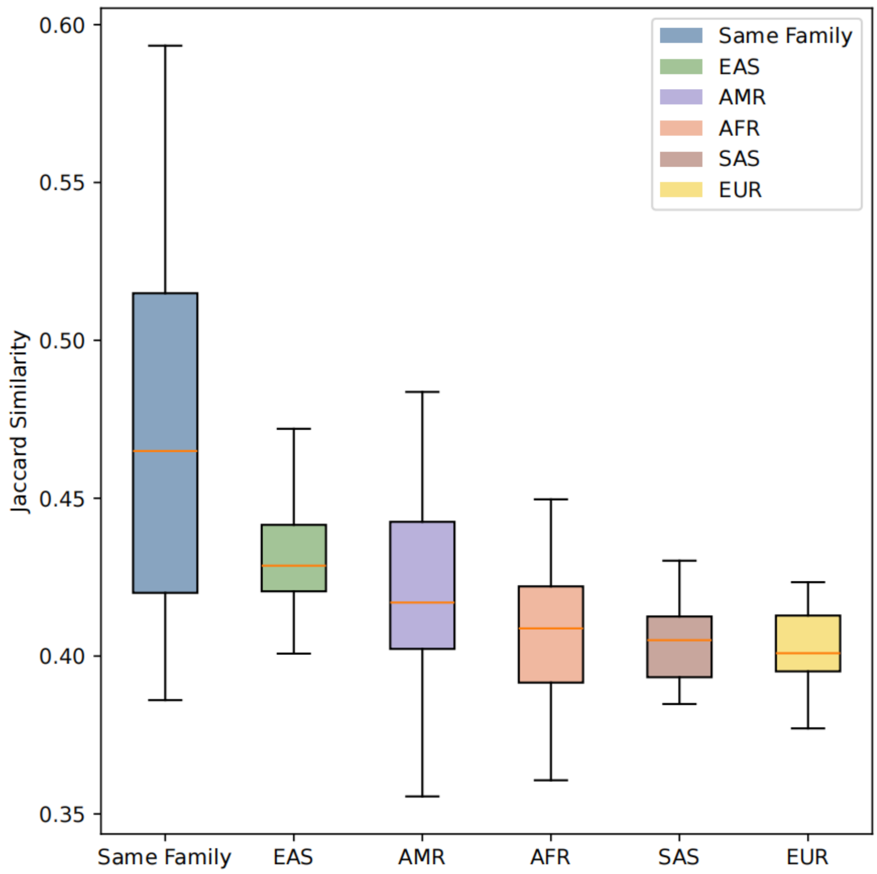


Supplementary Fig. 19 Boxplots of pairwise Jaccard similarities of novel STR variants uniquely identified by FastSTR, grouped by whether individuals belong to the same family. Jaccard similarities are significantly higher within families compared to between different families.

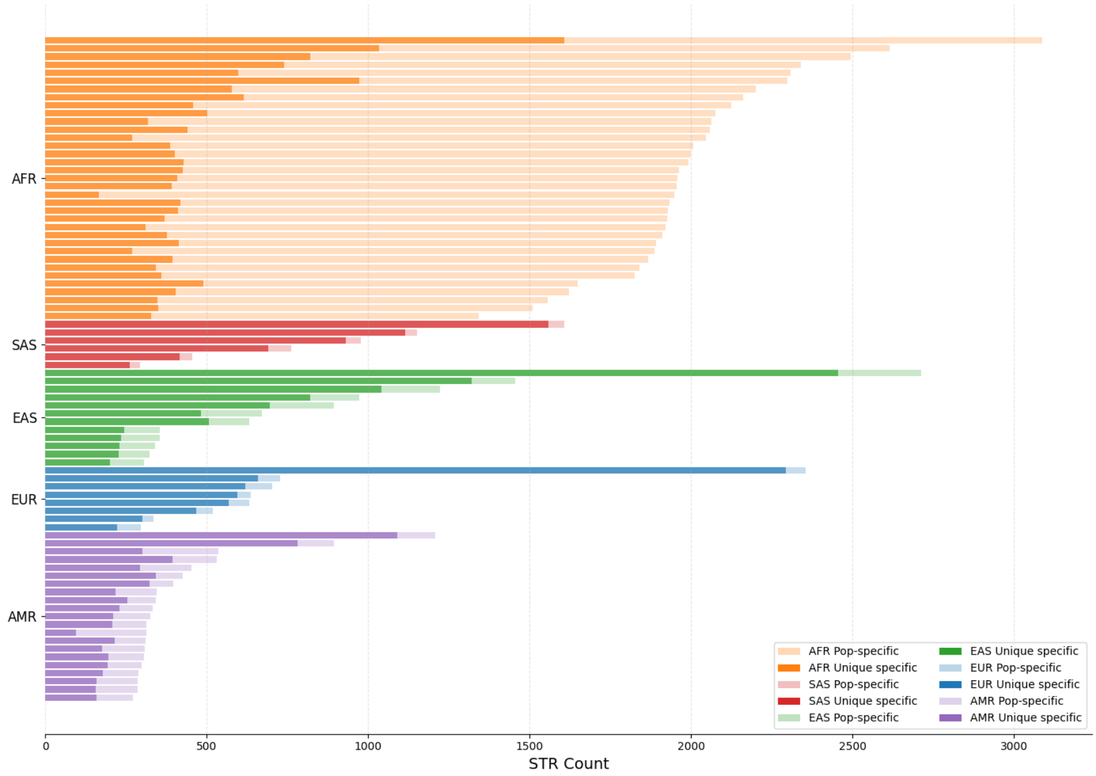


Supplementary Fig. 20 Number of super-population-specific and individual-specific novel STR variants uniquely identified by FastSTR across different populations. The African super-population exhibits the highest number of specific variants, whereas the American super-population shows the lowest.

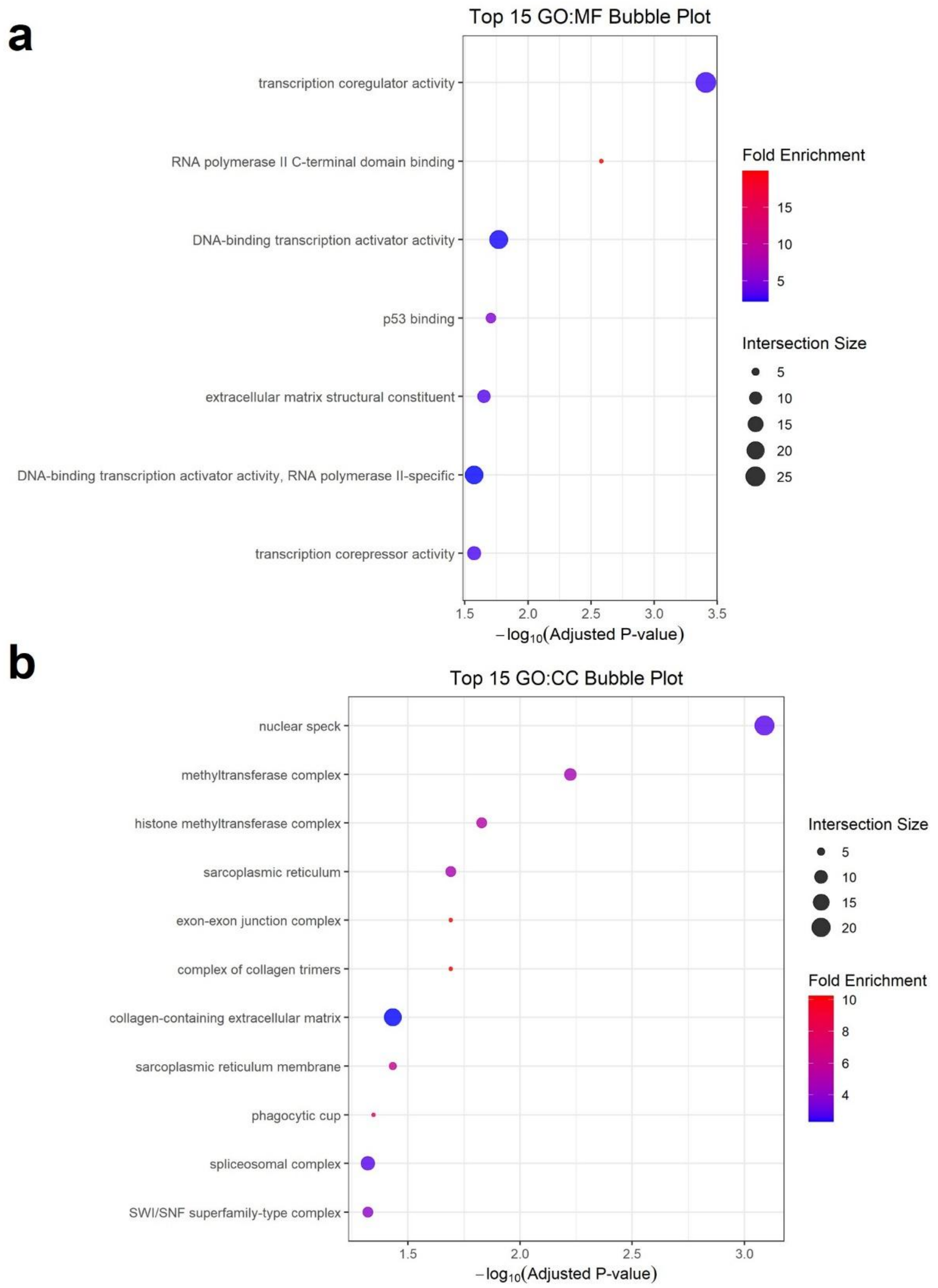


Supplementary Fig. 21 a. GO enrichment analysis (Molecular Function, MF) of genes affected by STR variants uniquely observed in lung cancer samples, located within coding sequences (CDS). b. GO enrichment analysis (Cellular Component, CC) of genes affected by STR variants uniquely observed in lung cancer samples within CDS regions.

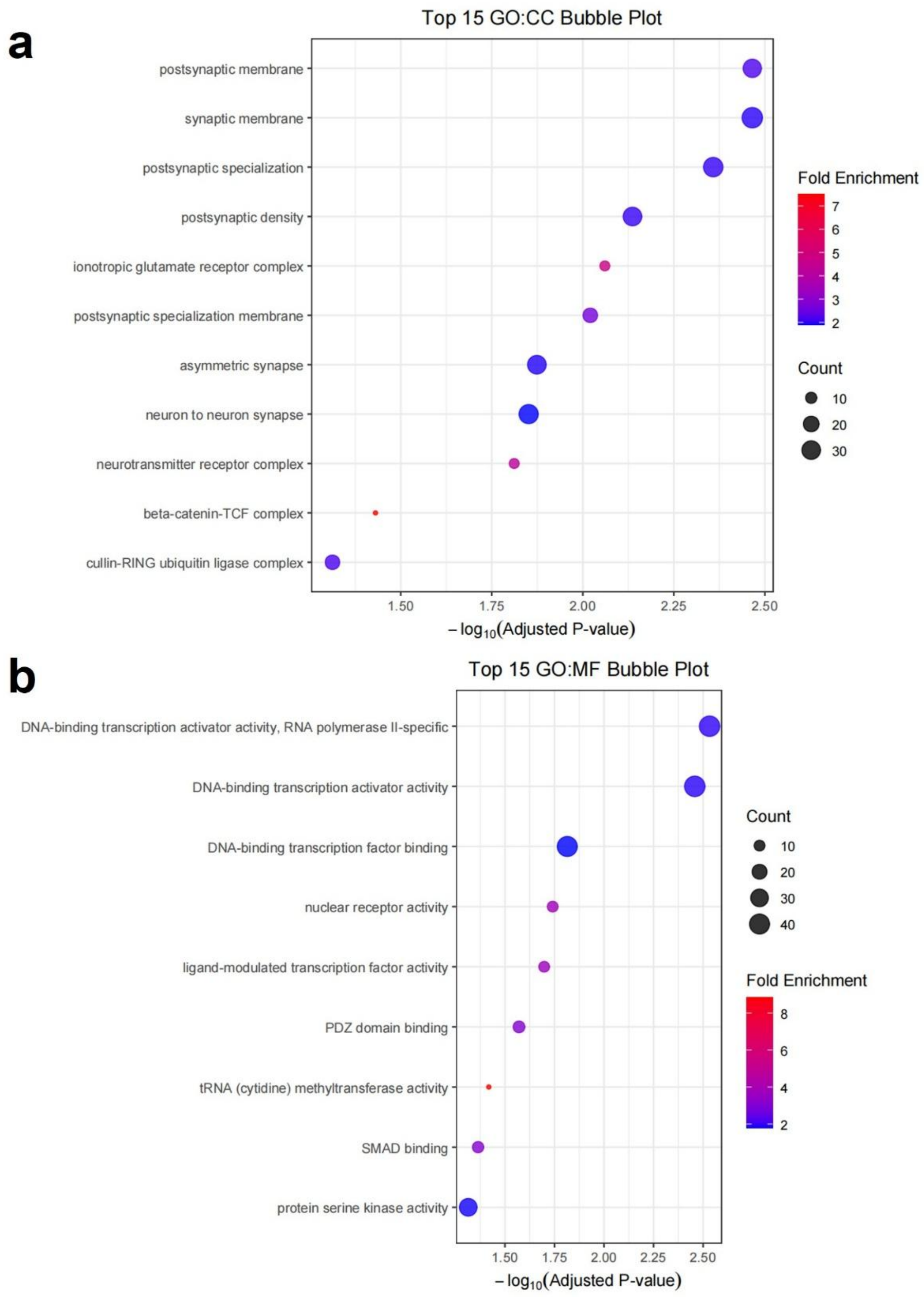


Supplementary Fig. 22 a. GO enrichment analysis (Molecular Function, MF) of genes affected by STR variants uniquely observed in lung cancer samples, located within untranslated regions (UTRs). b. GO enrichment analysis (Cellular Component, CC) of genes affected by STR variants uniquely observed in lung cancer samples within UTR regions.
